# Seed treatment with chitosan synergizes plant growth promoting ability of *Pseudomonas aeruginosa*-P17 in sorghum (*Sorhum bicolor* L.)

**DOI:** 10.1101/601328

**Authors:** G. Praveen Kumar, Suseelendra Desai, Bruno M. Moerschbacher, Nour Eddine-El Gueddari

**Affiliations:** Division of Crops Sciences, Central Research Institute for Dryland Agriculture, Santoshnagar, Saidabad Post, Hyderabad-500 059, India; Institute of Plant Biology and Biotechnology, University of Muenster, Hindenburgplatz-55, D48143 Muenster, Germany

**Keywords:** *Pseudomonas* P17, Chitosans, DA 56%, Plant growth promotion, CCME

## Abstract

Inoculation of crop plants with PGPR has in a large number of investigations resulted in increased plant growth and yield both in the greenhouse and in the field. This plant growth promoting effect of bacteria could be due to net result of synergistic effect of various pgpr traits that they exert in the rhizosphere region of the plant. Four (04) bacterial strain of fluorescent Pseudomonas spp. viz. P1, P17, P22 and P28 were identified previously for their plant growth promoting nature and abiotic stress tolerance and selected further to assess their chitinolytic activity and growth promotion on sorghum in combination with chitosans of low and high degree of acetylation. It was found that P1 has no chitin degrading nature and rest of the three strains have this property. When studied for their ability to grow in presence of chitosans of DA 1.6, 11, 35 and 56% all the strains showed growth in presence of chitosans. Seed bacterization of sorghum seeds with 04 bacterial strains in the presence and absence of chitosans (both low and high DA) and assessment of plant growth promotion after 15 days of sowing showed that P17+DA 56% chitosan combination showed higher growth of seedlings in plant growth chamber with highest root length of 25.9 cm, highest shoot length of 32.1 cm and dry mass of 132.7 mg/ plant. In P17+DA 56% chitosan treated seedlings various defence enzymes and PR-proteins were found to be present in highest quantities as compared to P1 and un-inoculated controls. Since this strain showed highest growth promotion of sorghum seedlings chitin-chitosan modifying enzyme (CCME) of this strain was partially characterized using different proteomic tools and techniques. CCME of P17 had one active polypeptide with a Pi in the range of 3.0-4.0. The digestion pattern of acetylated and deacetylated chitosans showed that P17 enzyme has endochitinase activity. Substrate specificity assay showed that the enzyme had more specificity towards highly acetylated chitosans. Two dimensional PAGE and MS analysis of the protein revealed similarities of this enzyme with protein of Pseudomonas aeruginosa chitinase PA01 strain of GenBank. In conclusion, the study established the option of opening new possibilities for developing bacterial-chitosan (P17+DA 56% chitosan) product for plant growth promotion and induced systemic resistance in sorghum.

## 1. Introduction

Plant growth-promoting rhizobacteria (PGPR) have been studied extensively for promoting plant growth and for inducing systemic resistance as well. PGPR-mediated induced systemic resistance (ISR) has been shown to effectively suppress several fungal, bacterial and viral pathogens in a number of crops both in greenhouse and field trials (Kloepper et al. 2004). Inoculation of crop plants with PGPR has in a large number of investigations resulted in increased plant growth and yield both in the greenhouse and in the field. The beneficial effects of these bacteria have in most cases been related to their ability to mobilize nutrients to plants, produce plant growth hormones and/or antimicrobial substances and to protect growing roots from deleterious root microorganisms present in the rhizosphere (Kloepper 1991; Bashan et al. 2004; Ortmann and Moerschbacher, 2006; Suseelendra Desai et al. 2012; Praveen Kumar et al. 2012; Leo Daniel Amalraj et al. 2015). Metabolites with biocontrol properties have been reported from diverse members of the rhizosphere flora. However, those produced by fluorescent *Pseudomonas* spp. have received a better attention which could be due to their abundance, proven bio-efficacy against several plant pathogens (Dowling and Gara, 1994).

Chitin is widely distributed in nature and forms a major constituent of the shells of crustaceans (such as crabs and shrimps), the exoskeletons of insects, molluscs, and cell walls of higher fungi. Chitinolytic enzymes have been purified from many sources and their enzymatic activities have been investigated. Chitinases (EC 3.2.1.14) catalyze the hydrolytic degradation of chitin and chitinases are classified into family 18 and family 19 of glycosyl hydrolases on the basis of their amino acids sequences (Henrissat and Bairoch, 1993). Potential use of natural enemies as alternatives or supplements for integrated plant disease management has been addressed in many studies (Morica et al. 2001; Lucas-Garcia et al. 2004). Many species of bacteria are known to synthesize chitinases for the utilization of chitin as a source of carbon and nitrogen. Some chitinolytic bacteria have been shown to be potential biological control agents, both of the plant diseases caused by various phytopathogenic fungi and of insect pests, because fungal cell walls and insect exoskeletons both contain chitin as a major structural component (Chernin et al. 1997; Manjula et al. 2004, Radjacommare et al., 2004; Oliviera et al. 2012). The hydrolysis products of chitin, chitooligosaccharides, are of interest in several biological and biotechnological processes (Peter, 2002; Synowiecki and Al-Khateeb, 2003; Subha Narayan et al., 2015) including agriculture to reduce use of chemical fungicides, which are leading to environmental pollution. Hence, as an alternative strategy, there is a growing interest on the application of chitins and chitosans in agriculture. Chitosan used to control plant pathogens has been widely explored with more or less success depending on the pathosystem, the used derivatives, concentration, degree of deacetylation, viscosity, application method; and chitosan alone or in combination with others (Abdelbasset et al. 2010).

Chitinases have been characterized from diverse microbial strains (Bröker et al. 2006; Meenavalli et al. 2010). These enzymes have applications not only in agriculture but also in other diverse commercial sectors. Neiendam Nielson and Sørensen (1999) reported an array of endochitinases and chitobiases in *P. fluorescens* and Folders et al (2001) reported chiC gene in *P. aeruginosa*. Chitinolytic activity would be an additional attribute for plant growth promoting bacteria as the trait could be used for breakdown of chitin and chitosan into oligomers with antimicrobial or defense inducing potential. The chitinolytic activity can help to modulate the rhizosphere environment where several complex interactions are involved and thus could render rhizosphere engineering for plant health management.

Out of several *Pseudomonas* spp. strains screened, we identified potential PGP strains (Praveen Kumar et al. 2015) and among them some strains also possessed abiotic stress tolerance (Goteti et al. 2014). This made us to look at our PGP strains of *Pseudomonas* spp. also for chitin and chitosan modifying enzymes (CCME). Hence, in this study, four selected strains of *Pseudomonas* possessing PGP ability in sorghum and/or possessing abiotic stress tolerance were used to characterize chitin-chitosan modifying enzymes and test their ability to promote growth of sorghum in presence of chitosan.

## 2 Materials and Methods

### 2.1 Bacterial growth conditions

Bacterial strains namely P1, P17, P22 and P28 were isolated by serial dilution and spread plating of bulk soil on King’s B medium (King et al. 1954) from farmerś fields of rainfed agro-ecosystems of India. Their identity was confirmed by 16s rRNA gene sequencing and standard biochemical tests.

### 2.2 Crude enzyme extraction

Cells of four bacterial strains (P1, P17, P22 and P28) were grown in 100 ml of tryptone-soy broth in 150 ml Erlenmeyer flask at 28°C and 140 rpm for 26 h. Cells were separated by centrifugation at 10000 rpm for 10 min. Supernatant was dried in liquid nitrogen (−196°C) and the samples were freeze-dried under vaccum in a lyophilizer at −60°C and 0.08 MPa pressure till the samples were totally dried. Crude dried powder (10 mg) expected to contain enzymes was dissolved in 1 ml of 50 mM sodium acetate buffer (pH 5.2) and further used to characterize the enzymes. The sample was desalted by passing through PD-10 columns as per the instructions of the manufacturer (GE, healthcare).

### 2.3 Substrates, and screening for Chitin-chitosan modifying enzymes (CCME) screening

Chitosan substrates with varying degrees of acetylation (DA) and polymerization (DP) *viz*, 1.6%, 11%, 35% and 56% were prepared by dissolving 1 mg of corresponding substrates in 1 ml of 100 mM glacial acetic acid and incubated overnight on a vortex shaker. Properties of various chitosans used in this study are given in Table 1.

**Table 1:** Details of *Pseudomonas* strains, their species, crop and location of isolation (in India) used in the present experiment

| ***Pseudomonas***<br>**strain** | **Species** | **NCBI**<br>**Accession No.** | **Crop** | **Location** | **State** |
| --- | --- | --- | --- | --- | --- |
| **P1** | *P. putida* | Not submitted | Greengram | Hayathnagar | Andhra Pradesh |
| **P17** | *P. aeruginosa* | Not submitted | Bulk soil | Baribrahmana | Jammu & Kashmir |
| **P22** | *P. aeruginosa* | HQ697284 | Sorghum-CSV 15 | Hayathnagar | Andhra Pradesh |
| **P28** | *P. aeruginosa* | HQ697285 | Cotton | Warangal | Andhra Pradesh |

Screening for the presence of CCME was done by dot blot assay on 5×5 cm (ref). polyacrylamide gels consisting 1 mgml^−1^ corresponding substrates Three ul of crude enzyme extract was applied on polymerized gels and incubated overnight in moist chamber at 37°C. The gels were stained following day with calcoflour white solution for 5 min and washed twice with distilled water for 15 min each and observed under UV light in a gel documentation system for enzymatic activity.

### 2.4 Selection of bacterial isolates for plant growth study in combination with chitosans

*Pseudomonas* isolates *viz*, P1, P17, P22 and P28 which were found to have pronounced effect on plant growth from the screening experiments were selected to evaluate their ability to enhance the growth of sorghum in combination with low and high DA chitosans.

#### 2.4.1 Effect of chitosans on bacterial growth

An experiment was carried out to test the effect of chitosans of different DA (56% and 1.6%) on growth of selected *Pseudomonas* isolates. The cultures were grown overnight to a population of 2×10^6^ CFU. mL^−1^. In a 96 well ELISA plate, 10 uL of bacterial cultures was added. To this culture, solutions of different chitosans were added to reach a final concentration of 5-200 µg. mL^−1^. The final volume of reaction mixture was made up to 150 uL per well. The plate was incubated in a ELISA reader for 18 h and OD was measured at 600 nm for every 10 min interval.

#### 2.4.2 PGP experiment

Effect of strains of *Pseudomonas* sp. either alone or in combination with chitosans of 1.4% and 56% DA on plant growth promotion of sorghum was studied. Seeds of sorghum *cv*. CSV-15 were treated as described elsewhere. Solutions of chitosans of DA 1.4% and 56% were prepared separately with a final concentration of 100 ug. mL^−1^. Actively growing bacterial strains (P1, P17, P22 and P28 @ 10^10^ cfu/ml) were applied as slurry along with chitosans were used to treat 12 sorghum seeds and final bacterial population was about 10^7^ cfu/seed before inoculation. The experimental setup consisted of un-inoculated control, seeds treated with chitosans alone and bacteria alone and chitosan-bacterial combinations for both the chitosans separately. Each treatment had three cups with four seeds and thinning was done to three seeds 5 DAS. Cups were watered regularly and placed in a plant growth chamber with light duration of 16 h (22°C) and dark duration of 8 h (17°C). Light intensity in the chamber was 70 umol/ m^2^/ sec (measured by Hansatech-Quantitherm light meter). After 15 days the plants were observed for increase in root, shoot length and dry mass. Experiment was repeated thrice.

### 2.5 Estimation of defence related enzymes in plants

As chitosan treatment is known to enhance the defence system in plants 15 DAS the fresh seedlings were used to estimate the quantity of different defence enzymes and pathogenesis related proteins.

#### 2.5.1 Estimation of phenylalanine ammonia lyase (PAL EC 4.3.1.5) activity

Leaf samples (1g) were homogenized in liquid nitrogen and extracted in 3 mL of ice-cold 0.1 M sodium borate buffer, pH 7.0, containing 1.4 mM 2-mercaptoethanol and 0.1 g insoluble polyvinyl-pyrrolidone. The extract was centrifuged at 16000 *g* at 4°C for 15 min. The supernatant was used as the enzyme source. Activity of PAL was determined as the rate of conversion of L-phenylalanine to *trans*-cinnamic acid as described by Dickerson et al. (1984). A sample containing 0.4 mL of enzyme extract was incubated with 0.5 mL of 0.1 M borate buffer, pH 8.8, and 0.5 mL of 12 mM L-phenylalanine in the same buffer for 30 min at 30°C. The optical density was recorded at 290 nm and the amount of *trans*-cinnamic acid formed calculated using its extinction coefficient of 9630 M^−1^ (Dickerson et al., 1984). Enzyme activity was expressed as µmol *trans*-cinnamic acid. min^−1^. g^−1^ protein.

#### 2.5.2 Estimation of peroxidase (PO EC 1.11.1.7) activity

Leaf samples (1 g) were homogenized in liquid nitrogen and extracted in 2 mL 0.1 M phosphate buffer, pH 7.0, at 4°C. The homogenate was centrifuged at 16000 *g* at 4°C for 15 min and the supernatant used as the enzyme source. The reaction mixture consisted of 1.5 mL of 0.05 M pyrogallol, 0.5 mL of enzyme extract and 0.5 mL of 1% H_2_O_2_. The reaction mixture was incubated at room temperature (28 ± 2°C). The changes in O.D. at 420 nm were recorded at 30 s intervals for 3 min. The enzyme activity was expressed as ‘Kat’ (Hammerschmidt et al. 1982).

#### 2.5.3 Estimation of poly-phenol oxidase (PPO EC 1.14.18.1) activity

PPO activity was determined as described by Mayer et al. (1965). Leaf samples (1 g) were homogenized in liquid nitrogen and extracted in 2 mL 0.1 M sodium phosphate buffer (pH 6.5) and centrifuged at 16000 *g* for 15 min at 4°C. The supernatant was used as the enzyme source. The reaction mixture consisted of 200 µL of the enzyme extract and 1.5 mL of 0.1 M sodium phosphate buffer (pH 6.5). To start the reaction, 200 µL of 0.01 M catechol was added and the change in O.D. was recorded at 30 s interval up to 3 min at 495 nm. The enzyme activity was expressed as ‘Kat’.

#### 2.5.4 Estimation of phenolics

Leaf samples (1 g) were homogenized in liquid nitrogen and extracted in 10 mL of 80% (v/v) methanol and agitated for 15 min at 70°C (Zieslin and Ben-Zaken, 1993). One mL of the methanolic extract was added to 5 mL of distilled water and 250 mL of Folin-Ciocalteau reagent (1 N) and the solution was kept at 25°C/ 3 min. Then 1 mL of 20% Na_2_CO_3_ was added and boiled in water bath for 1 min and the intensity of the developed blue colour was measured at 725 nm. Catechol was used as the standard. The amount of phenolics was expressed as ug catechol. mg^−1^ FW.

#### 2.5.5 Estimation of chitinase (EC 3.2.1.14) activity

Leaf samples (1 g) were homogenized in liquid nitrogen and extracted in 3 mL of 50 mM sodium acetate buffer of pH 5.2 and centrifuged at 16000 *g* for 15 min at 4°C. The supernatant was used as the enzyme source. The enzyme was incubated with DA 56% substrate and reducing ends were measured after incubation as described by Imoto and Yagishita (1971) and specific activity was expressed as nKat. mg^−1^ protein.

#### 2.5.6 Estimation of β-1, 3-glucanase (EC 3.2.1.39) activity

Leaf samples (1 g) were homogenized in liquid nitrogen and extracted in 3 mL of 50 mM sodium acetate buffer of pH 5.2 and centrifuged at 16000 *g* for 15 min at 4°C. The supernatant was used as the enzyme source. The activity of β-1,3-glucanase was determined by measuring the release of reducing sugars by using laminarin as substrate and glucose as standard. The reaction mixture consisted of 0.25 mL of enzyme solution, 0.3 mL of 1M sodium acetate buffer (pH 5.3) and 0.5 mL of 4% laminarin (Pan et al., 1991). The reaction was carried out at 40°C for 2 h. The reaction was stopped by adding 375 µL of dinitrosalicylic acid and heating for 5 min in a boiling water bath, vortexed and its O.D. measured at 500 nm. Protein concentration was determined by the method of Bradford (1976). The specific activity of β-1,3-glucanase was expressed as µg glucose released min^−1^ g^−1^ protein.

### 2.6 Enzyme assay and characterization

To the crude extract of P17 CCME 20 ul, different chitosan substrates at a concentration of 2 mgml^−1^ were mixed and incubated at 37°C overnight. Later, samples were concentrated at 30°C for 1 h. The pellet was dissolved in three ul distilled water and spotted onto 5×5 cm silica gel plates (Merck) using automatic TLC sampler. Samples were resolved using 41.7 ml n-Butanol, 33.3 ml methanol, 16.7 ml 25% ammonia and 8.3 ml distilled water. The spots were identified by spraying a solution prepared using 0.8 ml aniline, 0.8 gm diphenylamine, 40 ml acetone, 6 ml 85% phosphoric acid. Similarly, chitin (A) and chitosan (D) dimer to hexamer oligos were also incubated with the enzyme of P17 to study the digestion pattern of oligomers. Chitosans of DAs (25 ul) were incubated with 15 ul of crude enzyme and reducing ends were measured (Imoto and Yagishita 1971).

#### 2.6.1 Zymography, Molecular Sieve Chromatography & MALDI-o-TOF

The presence and number of iso-forms of enzyme in the crude extract were detected by zymography following the method of (Trudel and Asselin 1989). The gels were prepared consisting 1 mgml^−1^ chitosan substrates. Electrophoresis was carried out at a constant voltage of 40 V for 2 h. After the run, the gels were washed with sodium acetate buffer containing 1% (v/v) Triton X100, followed by washing with 50 mM sodium acetate buffer. Then the gels were incubated overnight at 37°C in 50 mM sodium acetate buffer. Next day, gels were stained with calcoflour white for five min and washed with water twice, and observed under UV light. Iso-electric focusing was carried out as per the manufacturer’s instructions (Amersham Biosciences, Inc.). Post IEF run, the gel was overlaid on DA 56% chitosan containing polyacrylamide gel and the iso-electric point of the enzyme was determined.

The average molecular weight and polydispersity index of the enzyme digested products were measured by gel permeation chromatography (Novema columns from PSS 30Å, 3000Å and guard column; 8 mm). Crude enzyme (50 ul) and DA 56% chitosan (950 ul) mixture was run on this column for 10 h to monitor the release of oligomers. Light intensity measurements were derived following the classical Rayleigh-Debye equation allowing us to deduce M̄_w_. The samples were eluted using degassed 0.2 M acetic acid/0.15 M ammonium acetate buffer (pH = 4.5). For accurate concentration measurements the refractive index increments (d*n*/d*c*) were determined independently for each sample using a differential refractometer (BI-DNDCW, Brookhaven Instruments Corporation, NYC, USA) in the same solvent, measured at 632.8 nm wave length.

P17 enzyme (20 ul) and DA 56% chitosan substrate (20 ul) mixture was incubated for 24 h and then first separated silica gel plates as described above and the spots were overlaid with glycerol (matrix) and the size and composition of the released oligomers was recorded using matrix assisted laser desorption/ ionization orthogonal time of flight (MALDI-TOF) procedure.

#### 2.6.2 2D-PAGE and protein identification by MS analysis of proteins

To identify the chitinase of P17, two dimensional polyacrylamide gel electrophoresis was performed (ref….). First dimension IEF was carried out as described above on 7 cm IPG strips and second dimension was carried out on polyacrylamide gel with DA 56% substrate. After the run, gel was incubated and the spot was picked, tryptic-digested and used for sequencing by mass spectrometry as described by Terashima et al. (2010). ‘Bioworks’ (Thermo) software was used for analysis of sequence data.

## 3. Results

### 3.1 Bacterial strains selection

Strains of *Pseudomonas* spp. P1, P17, P22 and P28 were selected for the current experiments based on their plant growth promotion towards sorghum, pigeon pea crops and abiotic stress tolerance (Praveen Kumar et al. 2012) (Table 1).

### 3.2 Dot blot assay

Dot blot assay revealed that P1 crude protein extract did not shown any dark colored spots conforming the absence of CCME. However P17 and P22 could show mild spots on DA 1.6% gel and their intensity increased with higher DA chitosans. In case of P28 spots developed on DA 11% and glycol chitin used gels (Table 2).

**Table 2:**
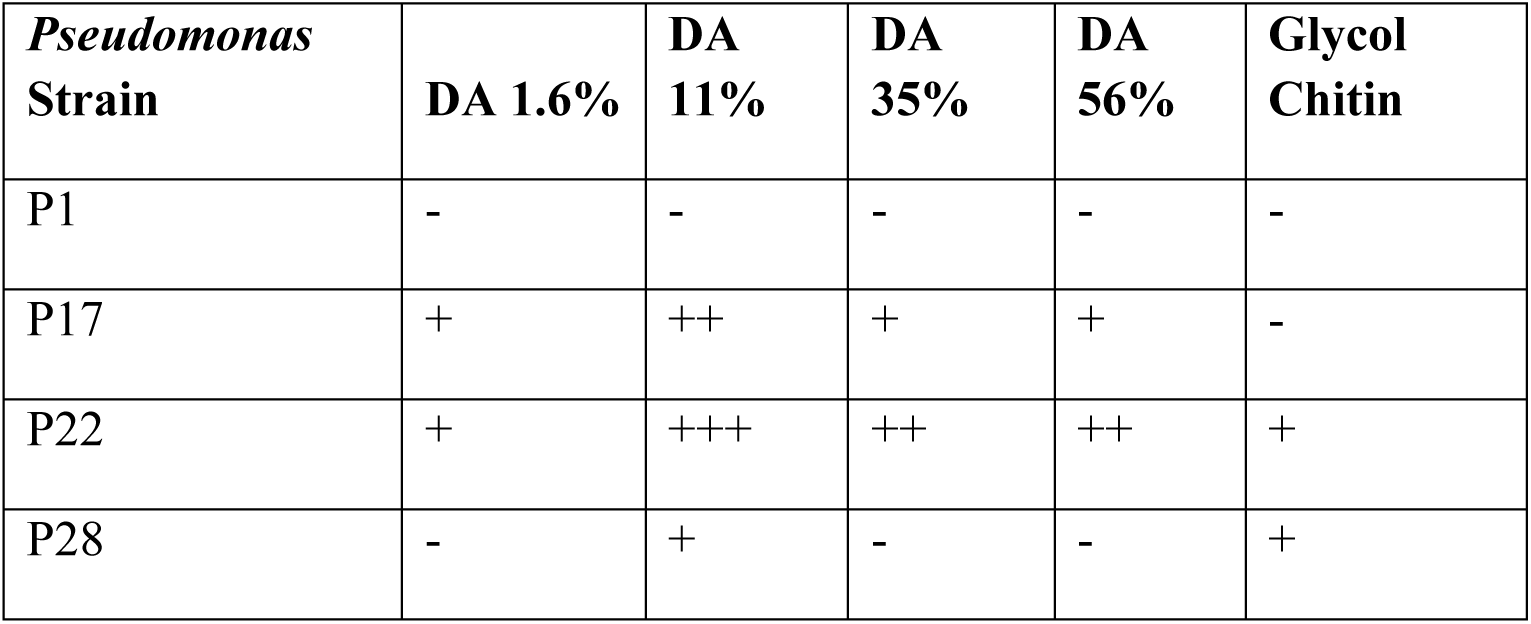
Production of chitinases/ chitosanases by *Pseudomonas* isolates

**Table 3:** Categories of chitosans and their properties

| Degree of Acetylation (%) | Degree of Polymerisation | Molecular Weight/Residue (g.mol<sup>-1</sup>) |
| --- | --- | --- |
| Parent chitosan |  |  |
| 1.5 | 2695 | 161.63 |
| High DP |  |  |
| 1.6 | 766 | 161.67 |
| 11 | 737 | 165.62 |
| 35 | 902 | 173.44 |
| 56 | 1442 | 184.52 |

### 3.3 Bacterial strains for plant growth studies in combination with/ without chitosans

#### 3.3.1 Bacterial growth in presence of chitosans

All the test bacterial strains showed growth in presence of both DA 1.6% and 56% chitosans. When 1.6% chitosan was used higher growth was recorded with P28 strain followed by P22. P1 and P17 did not grow to higher extent like that of other two strains. In presence of 56% chitosan, higher growth was recorded. All the strains had the similar growth rate with high DA chitosan. All strains recorded higher growth in complete absence of chitosans (Fig 1).

**Fig 1:**
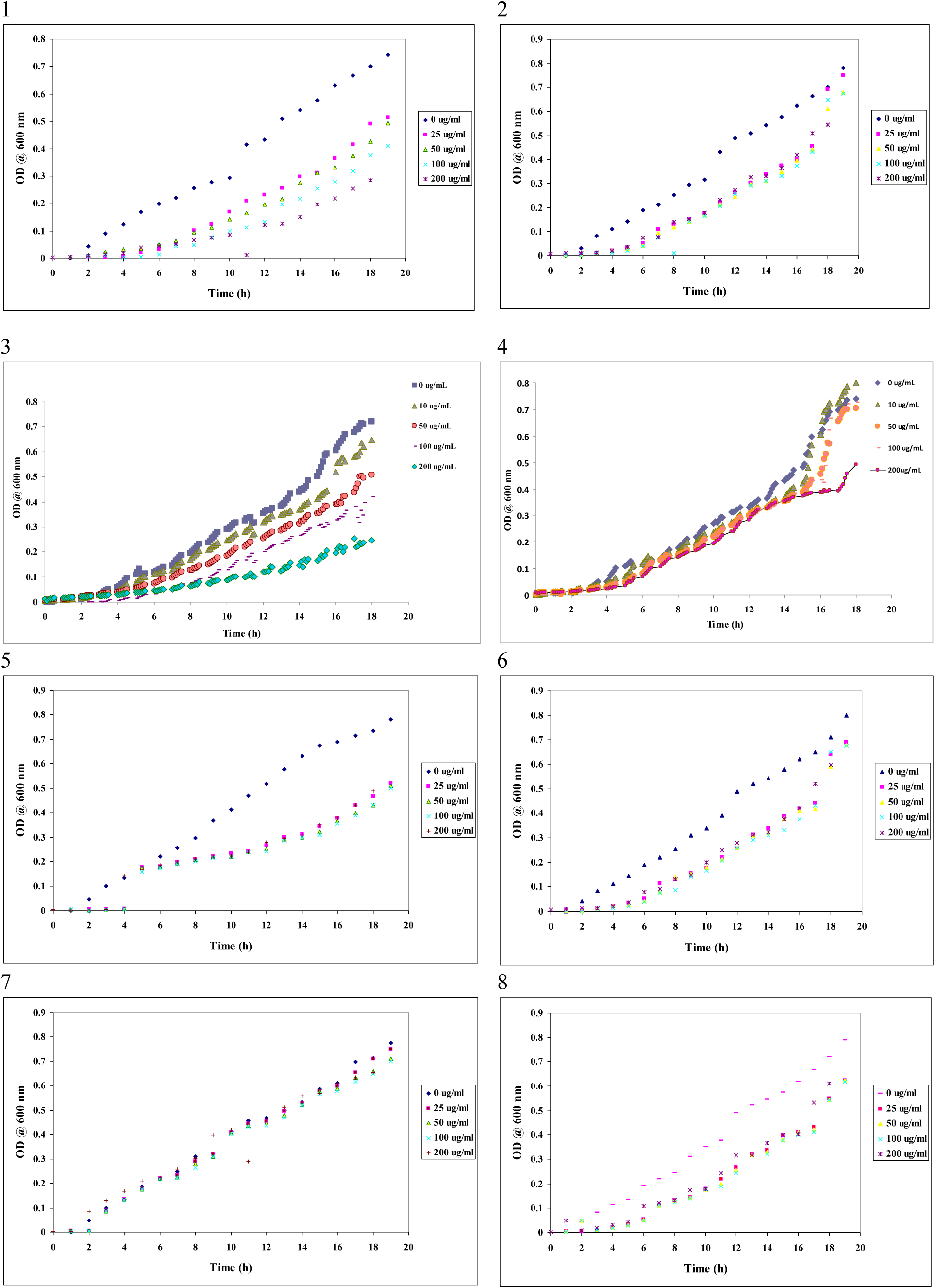
Effect of chitosans DA 1.6% and DA 56% on growth of P1 (1 & 2), P17 (3 & 4), P22 (5 & 6 and P28 (7 & 8) isolates, respectively.

#### 3.3.2 Plant growth promotion test

Using sorghum as a test plant, four bacterial isolates and two chitosans were tested either alone or in combination for their ability to promote plant growth. As compared to control, all the treatments promoted plant growth significantly. However, the combinations of *Pseudomonas* and chitosans outperformed rest of the treatments. The root length across treatments ranged from 19.4 to 25.9 cm. P17 coupled with DA 56% chitosan showed the highest root length of 25.9 cm followed by P1+DA 56% and P17 which were 24.6 and 23.4 cm, respectively. The treatments comprising P1, P22 and P28 or P1, P17 and P22 coupled with 1.4% DA were at par showing root length ranging from 20.8 to 21.2 cm (Table 4). The shoot length of the seedlings ranged from 20.0 to 32.1 cm across the treatments. Highest shoot length was observed in P17+DA 56% treatment which was statistically superior to all other treatments (Fig 2). The next best treatments were P1+DA 56% (28.1 cm) and P28+DA 56%. DA 1.4% treated plants showed shoot length of 22.1 cm and control plants showed only 20.0 cm (Fig 2).

**Fig 2:**
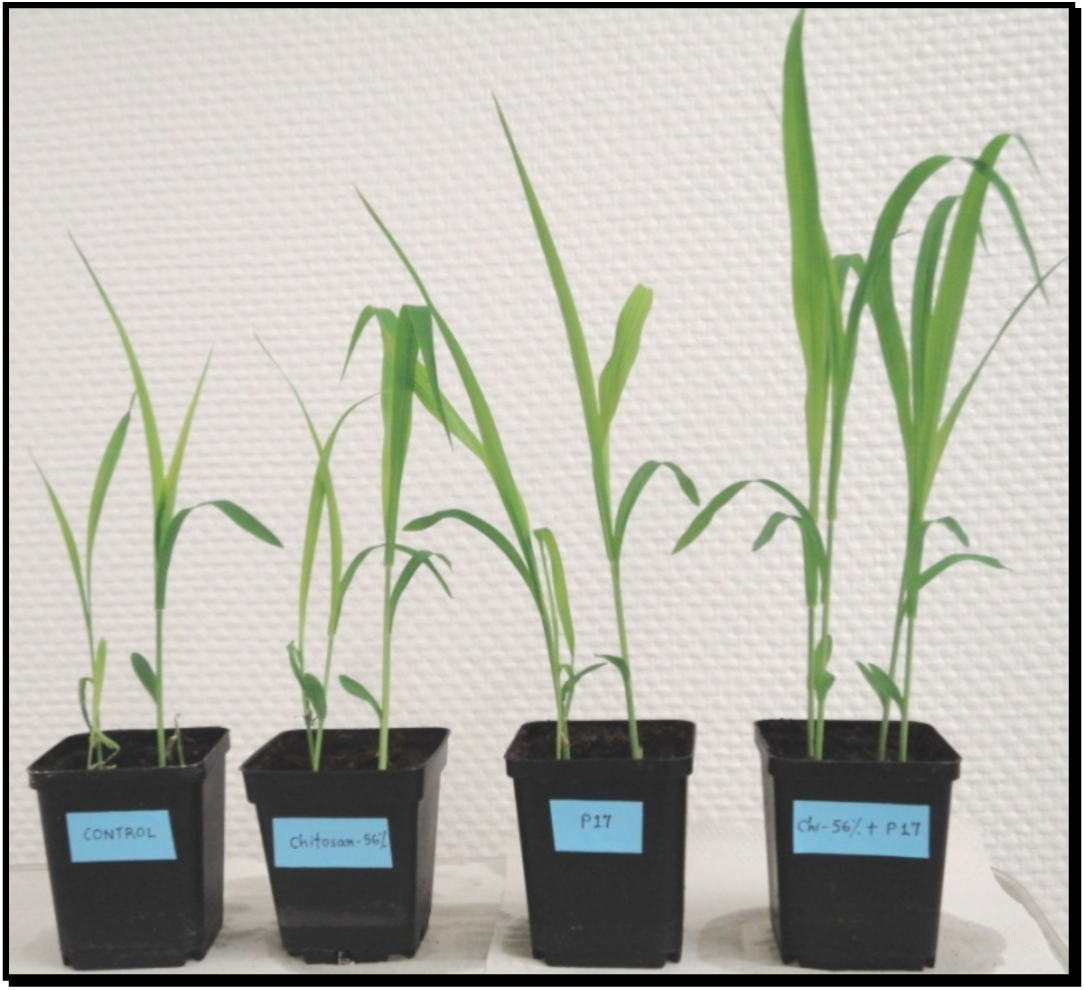
Plant growth promotion by *P. aeruginosa* P17 strain seed bacterization in combination with chitosan DA 56% on sorghum (15 DAS)

**Table 4:**
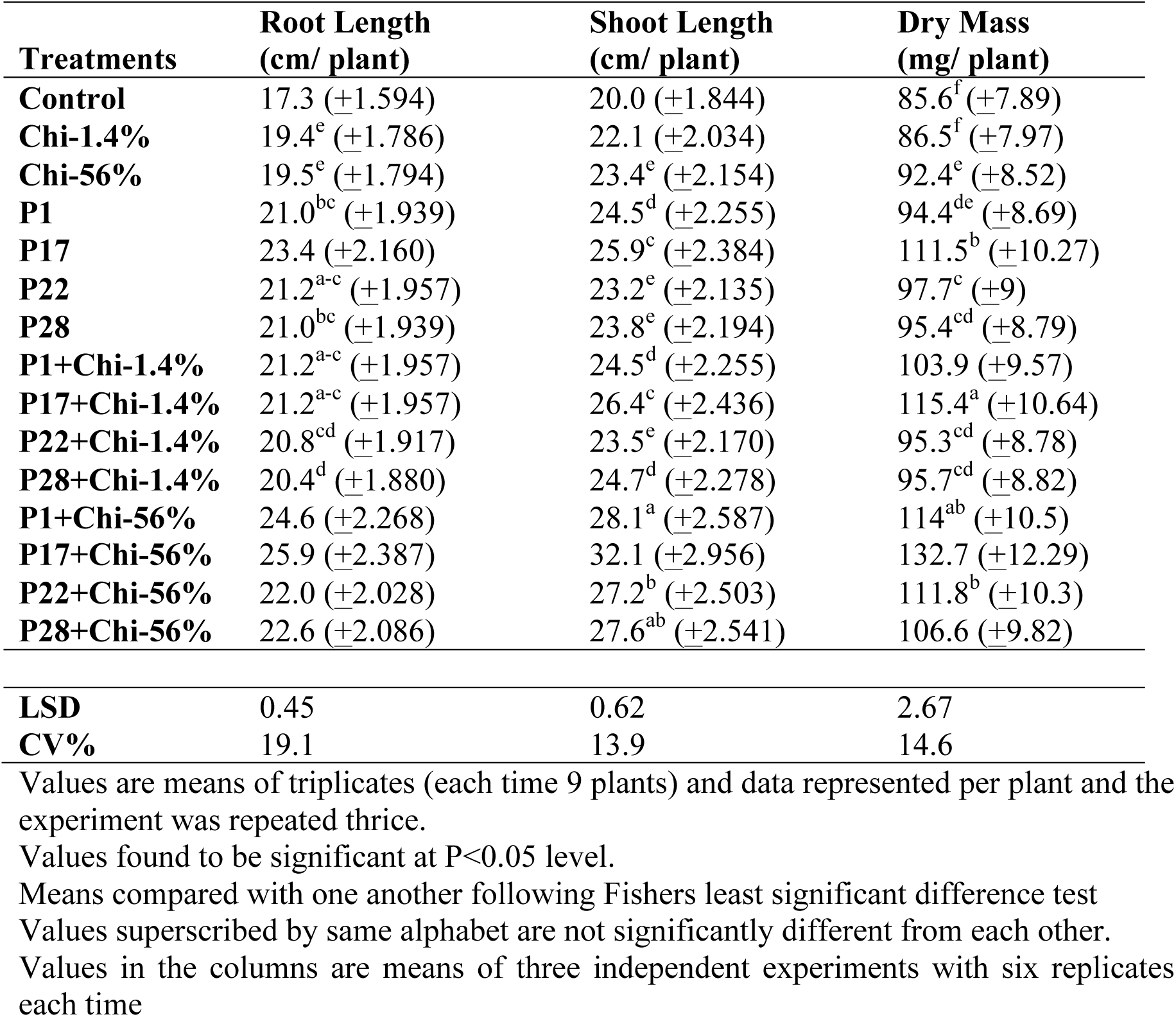
Changes in root, shoot length and dry mass of sorghum plants when inoculated with four bacterial strains and low, high DA chitosans (15 DAS)

Of all the treatments P17+chi-56% treated plants recorded higher dry mass of 132.7 mg followed by P17+DA 1.4% plants which showed 115.4 mg where as control plants showed only 85.6 mg. P17 in combination with DA 56% chitosan could enhance root, shoot length and dry mass by 50, 60 and 55%, respectively. Scanning electron micrograph of seeds after treatment showed the presence of P17 bacteria on their surface (Fig 3).

**Fig 3:**
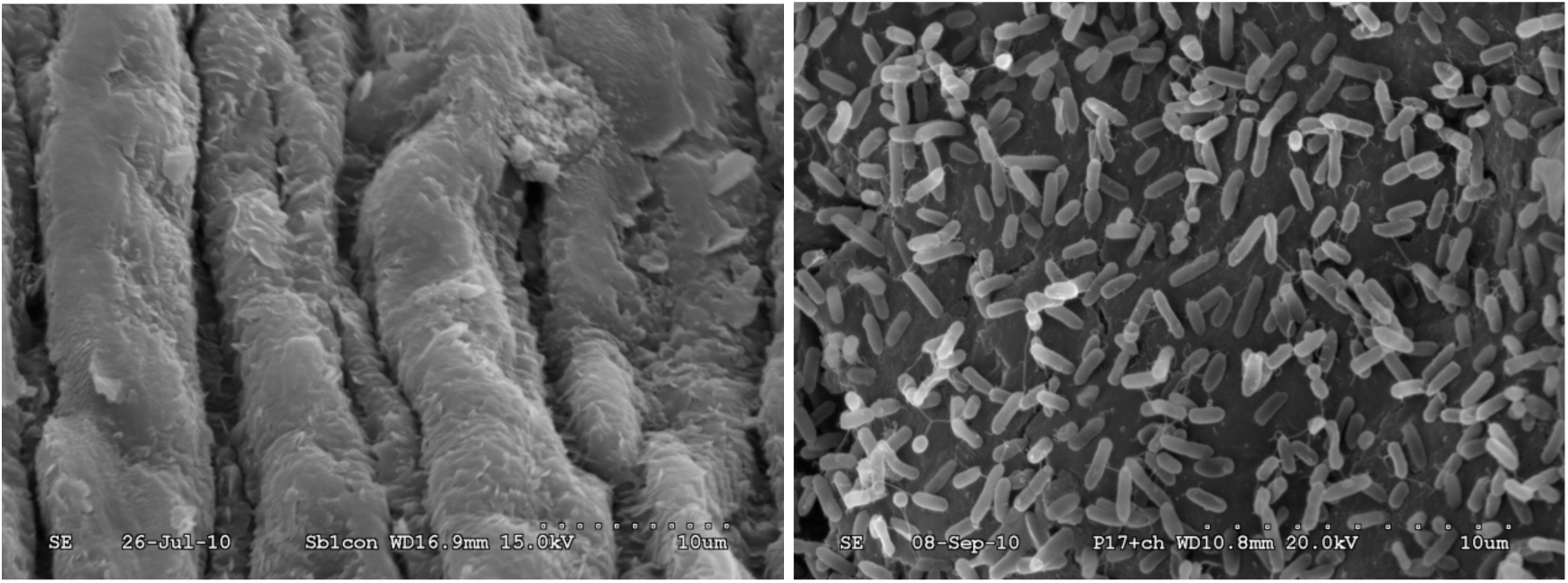
Scanning electron micrographs of sorghum seed showing the presence of chitosan and bacteria (right) and untreated control (left) at a magnification 5000X

#### 3.3.3 Defence enzymes in plants

##### PAL enzyme

Since, DA 56% chitosan is known to induce defence reactions in plants, different defense related enzymes were assessed in treated plants. Higher phenyl alanine ammonia lyase (PAL) activity was observed in plants treated with P22+DA 56% chitosan plants which was 0.05 umol cinnamic acid/ min/ gm, followed by DA 1.4% treatment and P17+DA 1.4% and P17+DA 56% treated plants which was 0.042 umol/ min/ gm which were on par. Control plants showed the lowest PAL activity (0.0026 umol/ min/ gm) compared to all other treatments (Fig 4).

**Fig 4:**
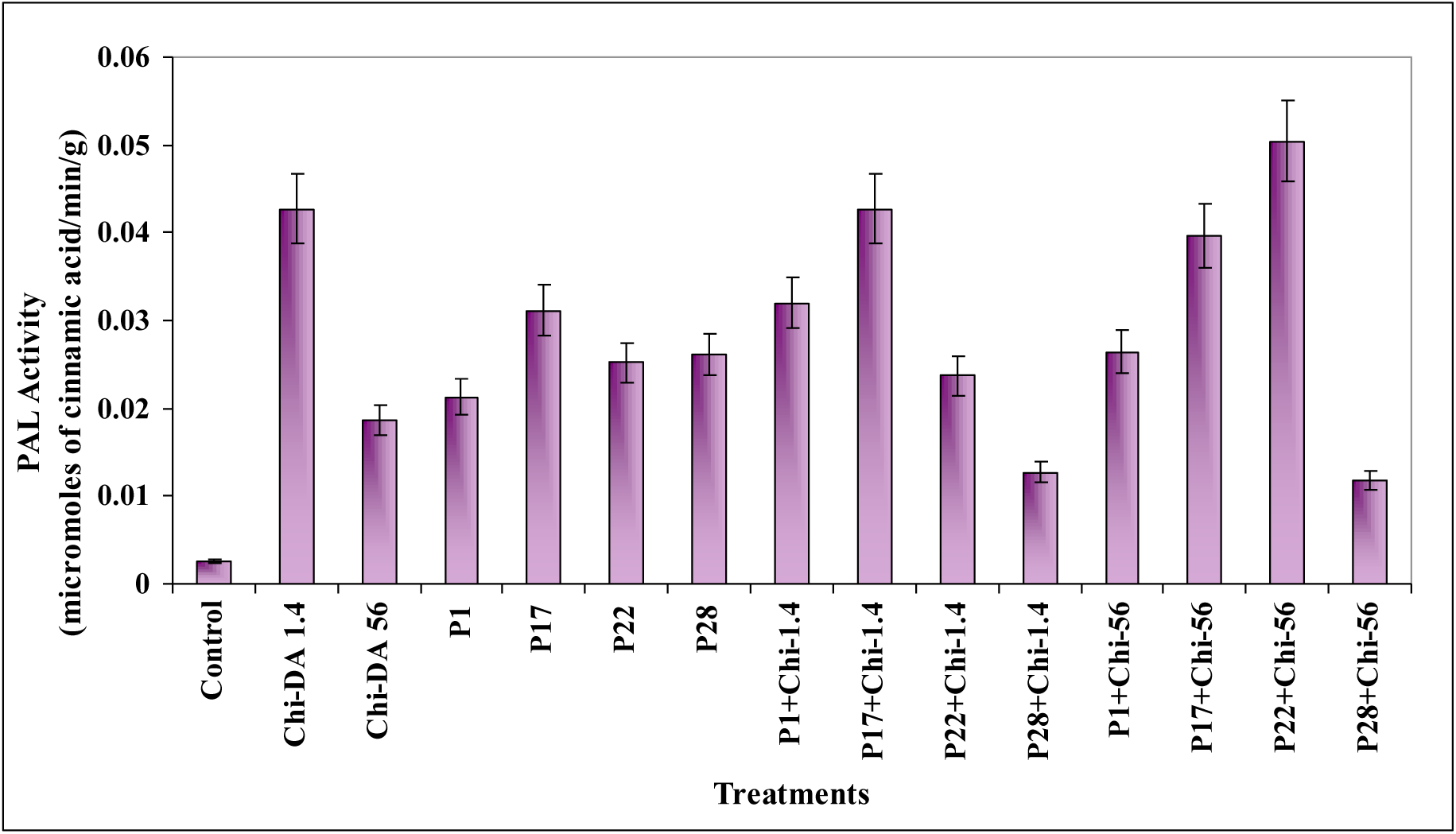
Effect of seed bacterization in combination with chitosans of different DA on induction of phenyl alanine ammonialyase in sorghum leaves (15 DAS). Values are means of three replicates

##### POX enzyme

For peroxidases (POX) control plants recorded lower activity of 0.0098 Kat. Whereas, P17+DA 56% treated plants (0.068 Kat) showed highest activity of POX followed by P1 and P17 (0.054 Kat) which were on par (Fig 5).

**Fig 5:**
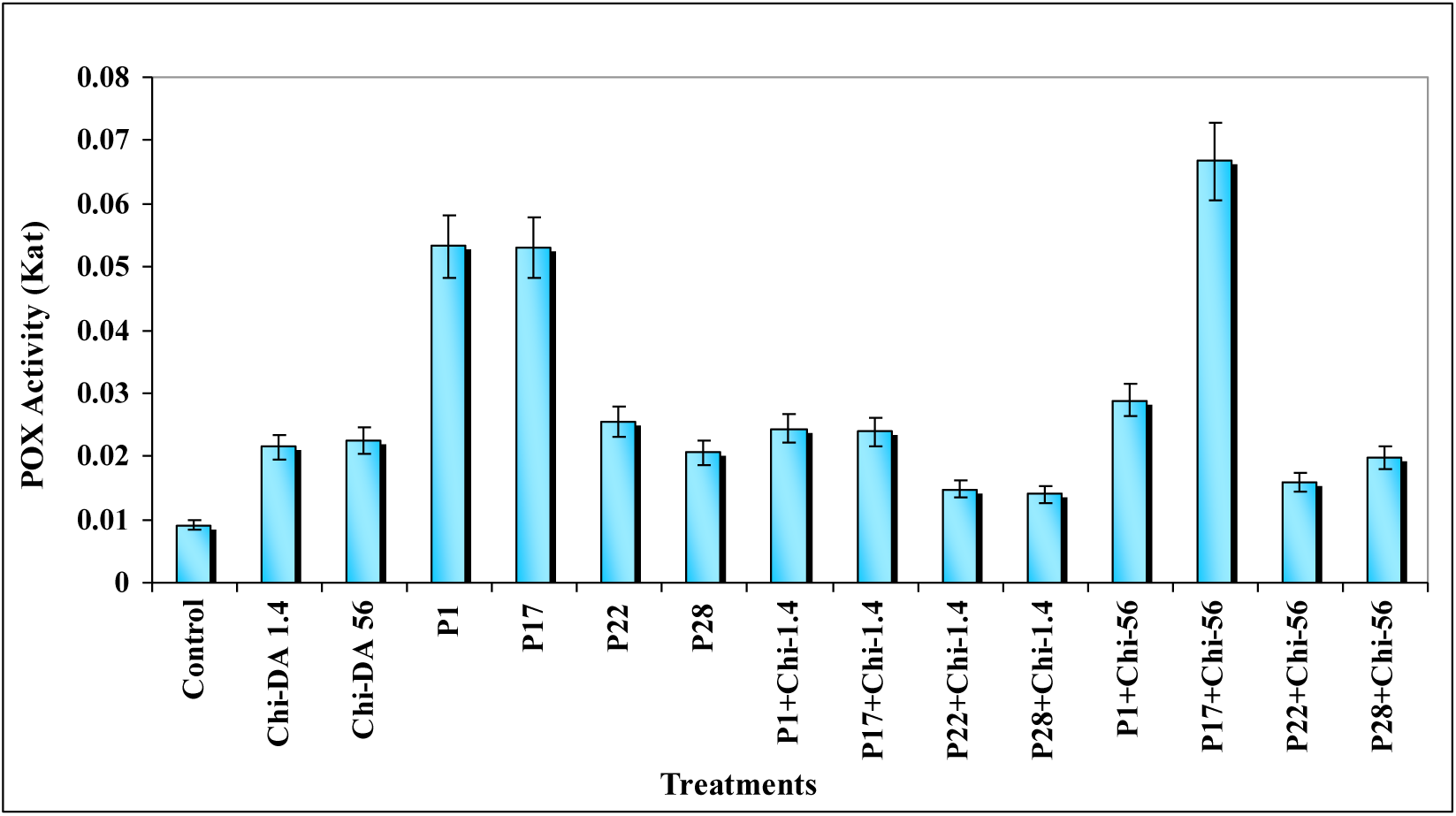
Effect of seed bacterization in combination with chitosans of different DA on induction of peroxidase in sorghum leaves (15 DAS). Values are means of three replicates

##### PPO enzyme

Polyphenol oxidase (PPO) activity was highest in case of P17+56% plants (0.0032 Kat). P17+1.4% plants showed a PPO activity of 0.0026 Kat followed by plants treated with P1 and P28 in combination with 56% which were on par (Fig 6). Individual chitosan and bacterial treated plants showed activity in the range of 0.0018-0.0019 Kat.

**Fig 6:**
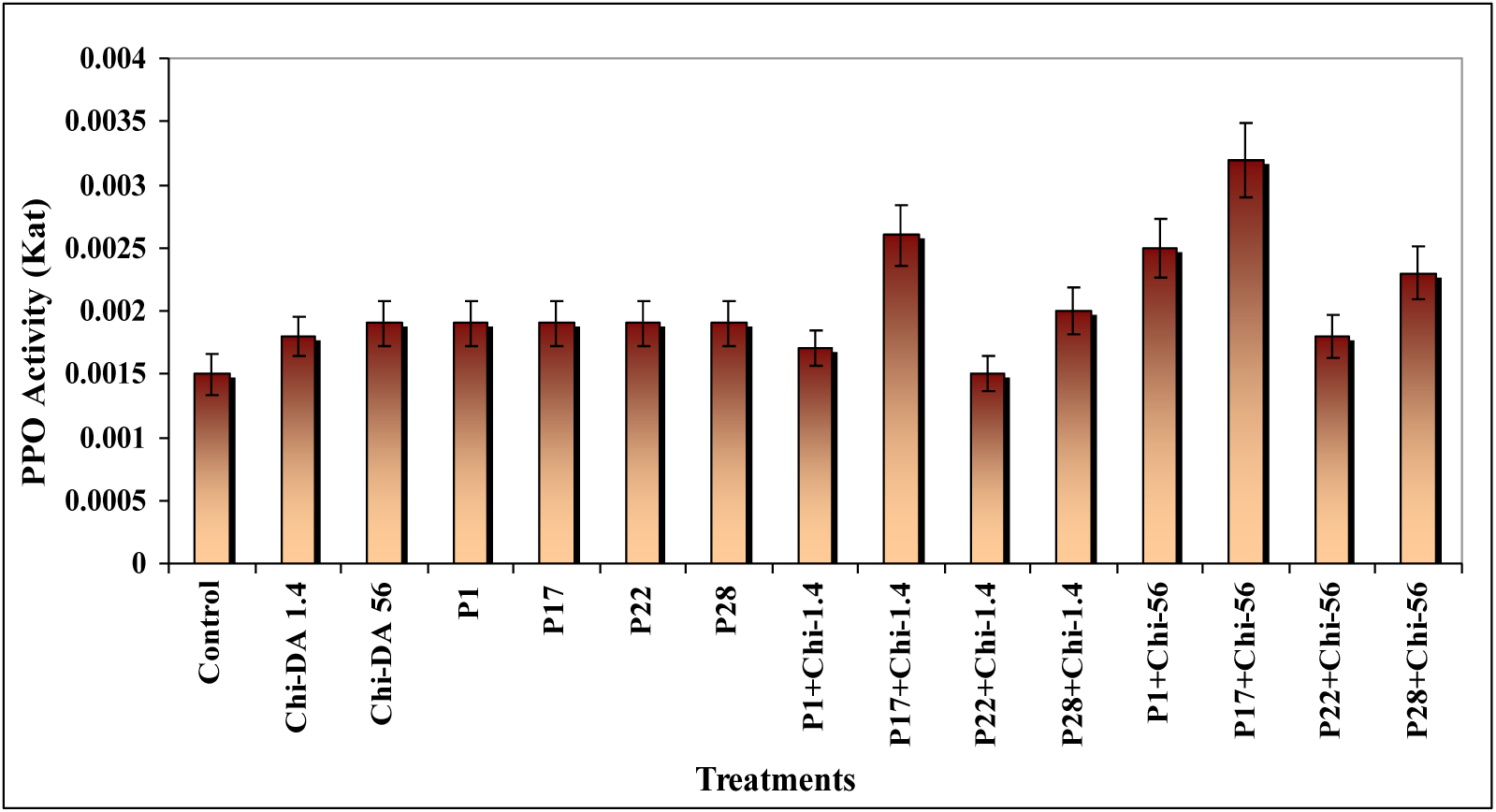
Effect of seed bacterization in combination with chitosans of different DA on induction of poly phenol oxidase in sorghum leaves (15 DAS). Values are means of three replicates

##### Phenolics content

When total phenolic content of the plants was analysed, all the treatments had more or less the same quantity of accumulated phenols where as in P17+56% chitosan (89.2 ug/ gm tissue) treated plants their quantity rapidly increased and control plants recorded only 22 ug of phenol/ gm of tissue (Fig 7).

**Fig 7:**
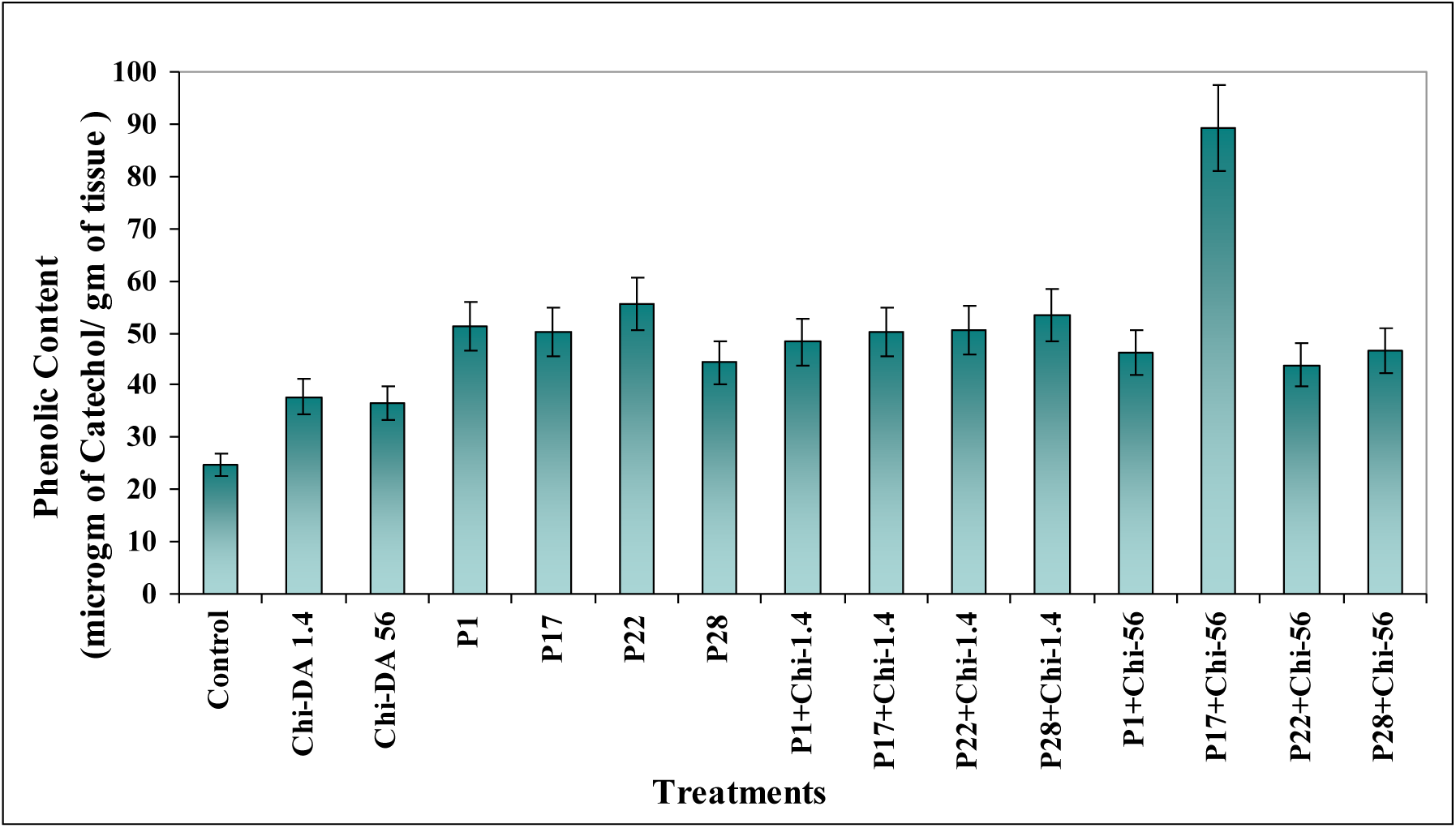
Effect of seed bacterization in combination with chitosans of different DA on induction of phenols in sorghum leaves (15 DAS). Values are means of three replicates

##### PR proteins

In case of pathogenesis related proteins, higher chitinase activity was observed in plants treated with P17+56% combination which was 3.32 nKat/ mg protein followed by P28+56% with 2.53 nKat/ mg protein activity where as control plants showed an activity of 0.403 nKat/ mg protein (Fig 8). Estimation of glucanase showed that, P17+56% plants recorded highest activity of 4500 nmoles/ min/ gm followed by P1+56% which was 3547 nmoles/ min/ gm activity whereas, untreated plants showed only 1100 nmoles/ min/ gm glucanase activity (Fig 9).

**Fig 8:**
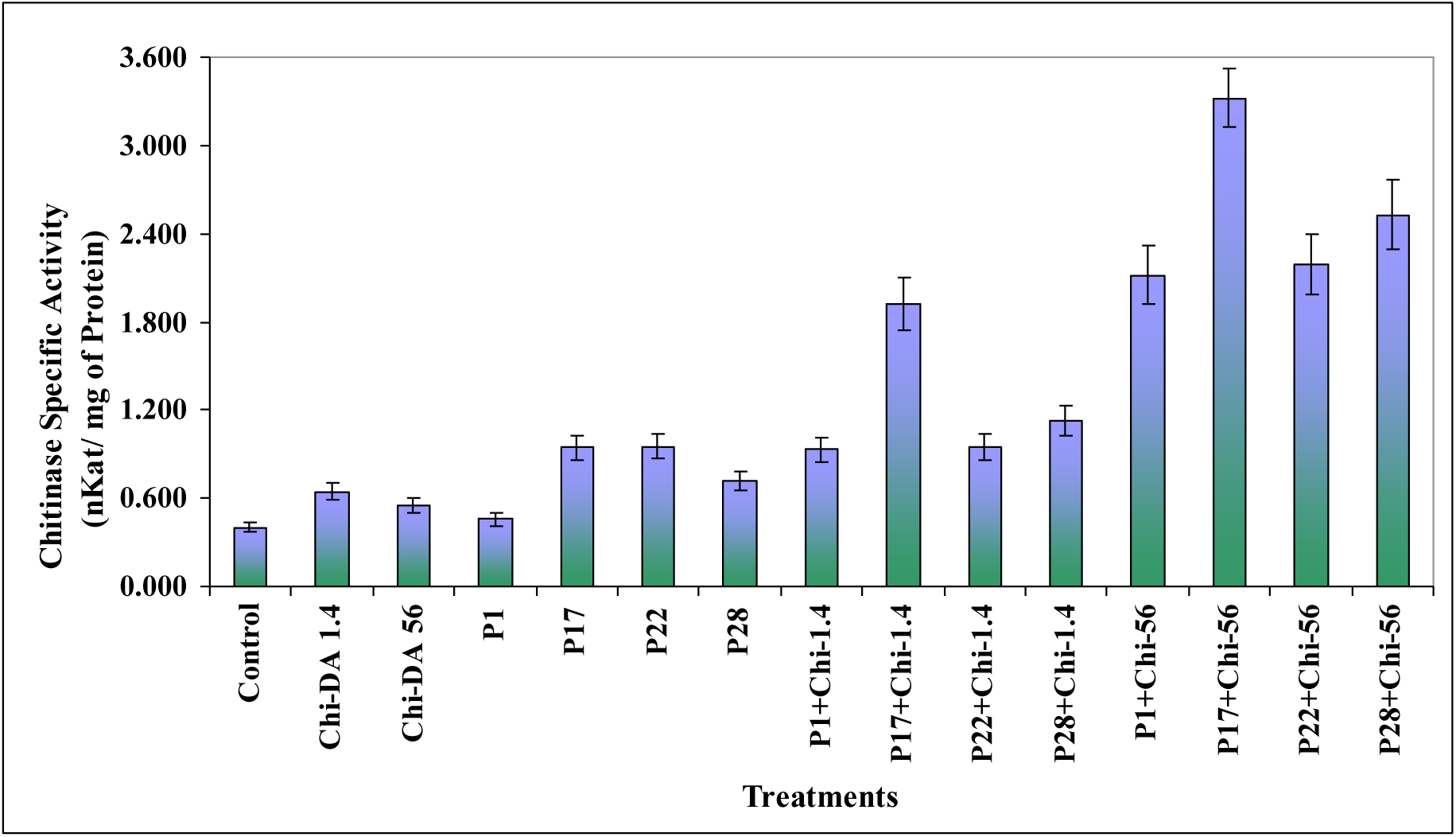
Chitinase specific activity in sorghum leaves as influenced by seed bacterization in combination with different chitosans (15 DAS). Values are means of three replicates

**Fig 9:**
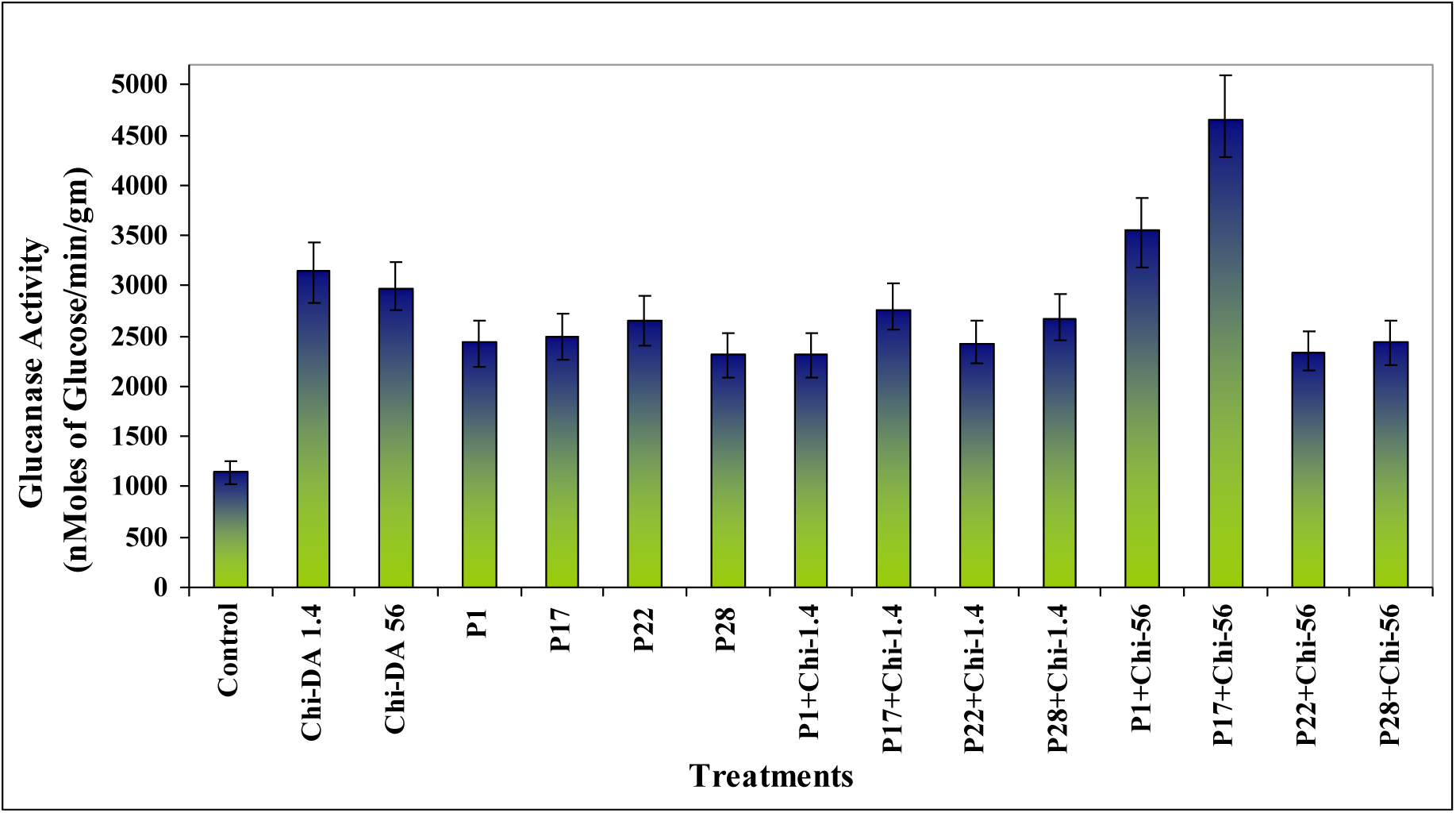
β-1, 3 Glucanase specific activity in sorghum leaves as influenced by seed bacterization in combination with different chitosans (15 DAS). Values are means of three replicates

#### 3.3.4 Enzyme activity & Characterization of P17 CCME

Based on the above results conducted with sorghum plants and P17 and chitosans since P17 found to have profound effect on plant growth in combination with Chitosan-DA 56% *Pseudomonas* P17 strain CCME was further characterized for revealing its features. Enzyme activity was quantitated with chitosan DA 56% as substrate. One unit of enzyme activity was defined as the amount of enzyme that liberated one umole of reducing sugar from the substrate per min with glucosamine as substrate. Total concentration of protein in the crude extract was 5.23 mgml^−1^. Specific activity of P17 CCME was calculated as 12.2 nKatmg^−1^.

##### Detection of isoforms by zymogprahy and iso-electric point

Semi-native gel assay for chitin-chitosan modifying enzymes showed that P17 isolate has one active protein for the hydrolysis of chitin/chitosans (Fig 10) and iso-electric focusing revealed that the Pi of the enzyme was between pH 3.0-4.0 with only one isoform (Fig 11).

**Fig 10:**
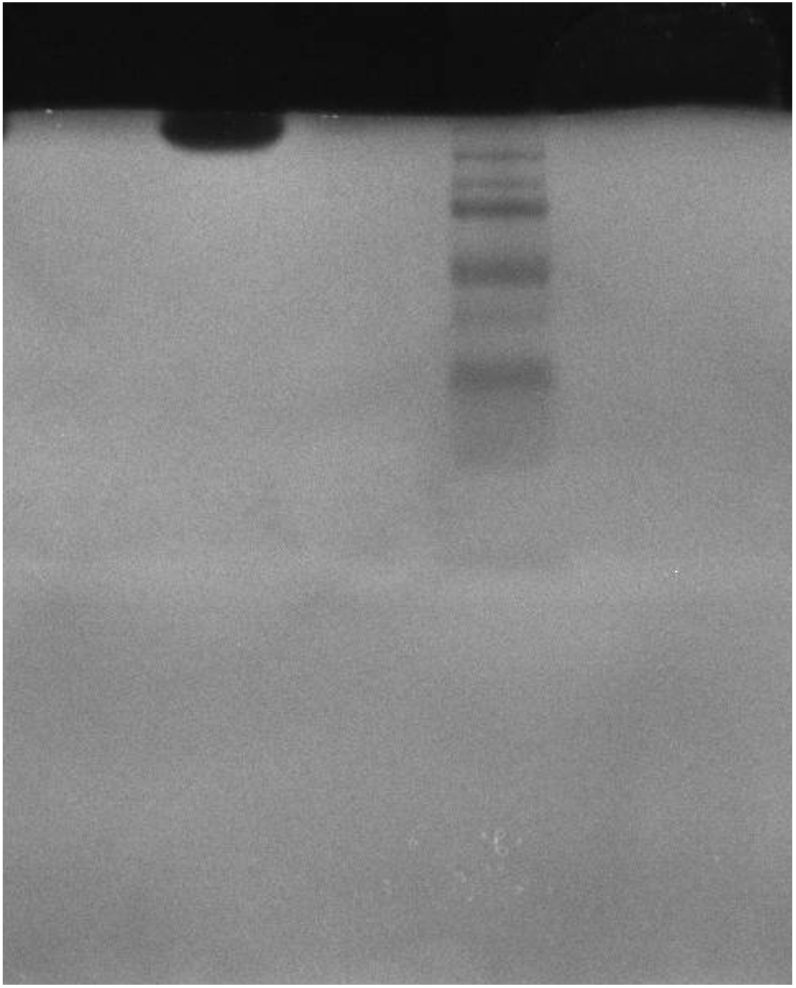
Semi-native PAGE gel showing the active protein of *Pseudomonas aeruginosa* (P17 CCME (Lane 1) and Molecular marker (Lane 2)

**Fig 11:**
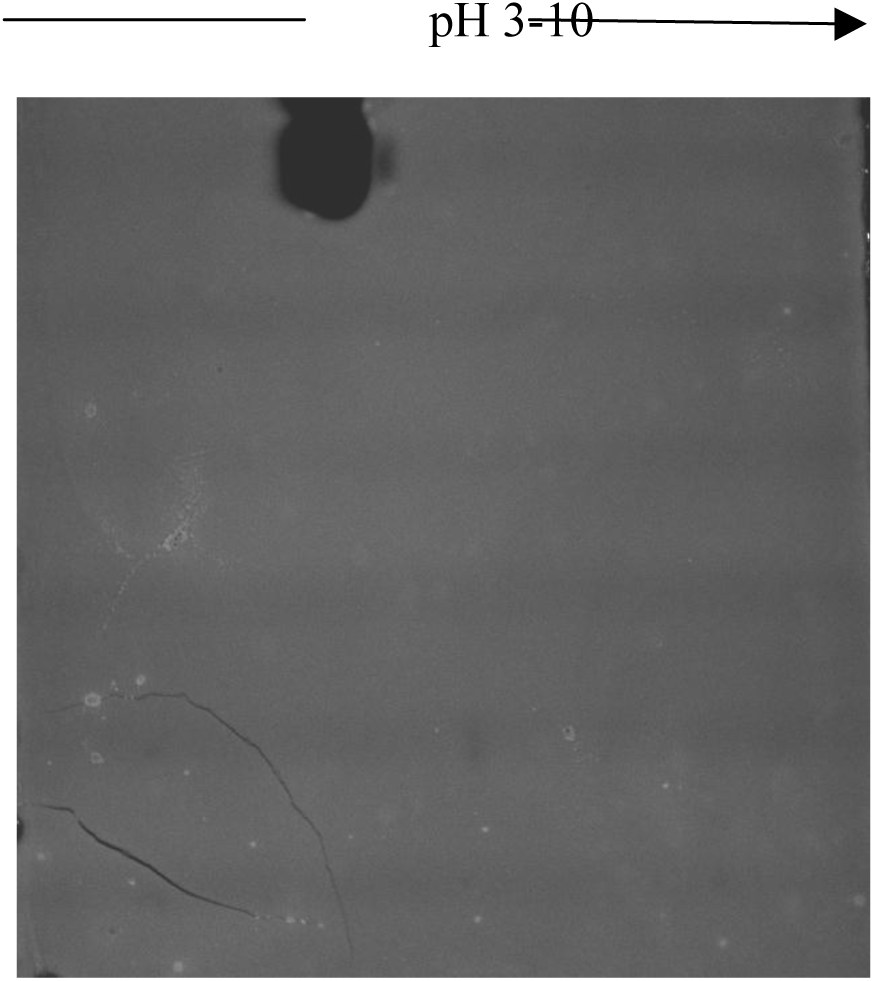
Iso-electric focussing overlay gel (with DA 56% chitosan) showing one isoform of *P. aeruginosa* P17 CCME

##### Substrate specificity

Reducing end assay was performed to assess the substrate specificity of the enzyme. Of the four chitosans of different DAs tested to characterize the hydrolytic properties of the CCME of P17 isolate, hydrolysis increased with the increase in DA. The number of micromoles of reducing ends released by the enzyme was high with chitosan DA 56% (0.11) and less with 1.6% DA chitosan (0.022). When DA 35% and 11% substrates were used, the reducing ends were 0.105 and 0.05 uMoles, respectively (Fig 4.30). Therefore, the substrate specificity of the P17 CCME in decreasing order was DA 56% > DA 35% > DA 11% > DA 1.6% suggesting that the CCME of P17 had high chitinase activity and low chitosanase activity (Fig 12).

**Fig 12:**
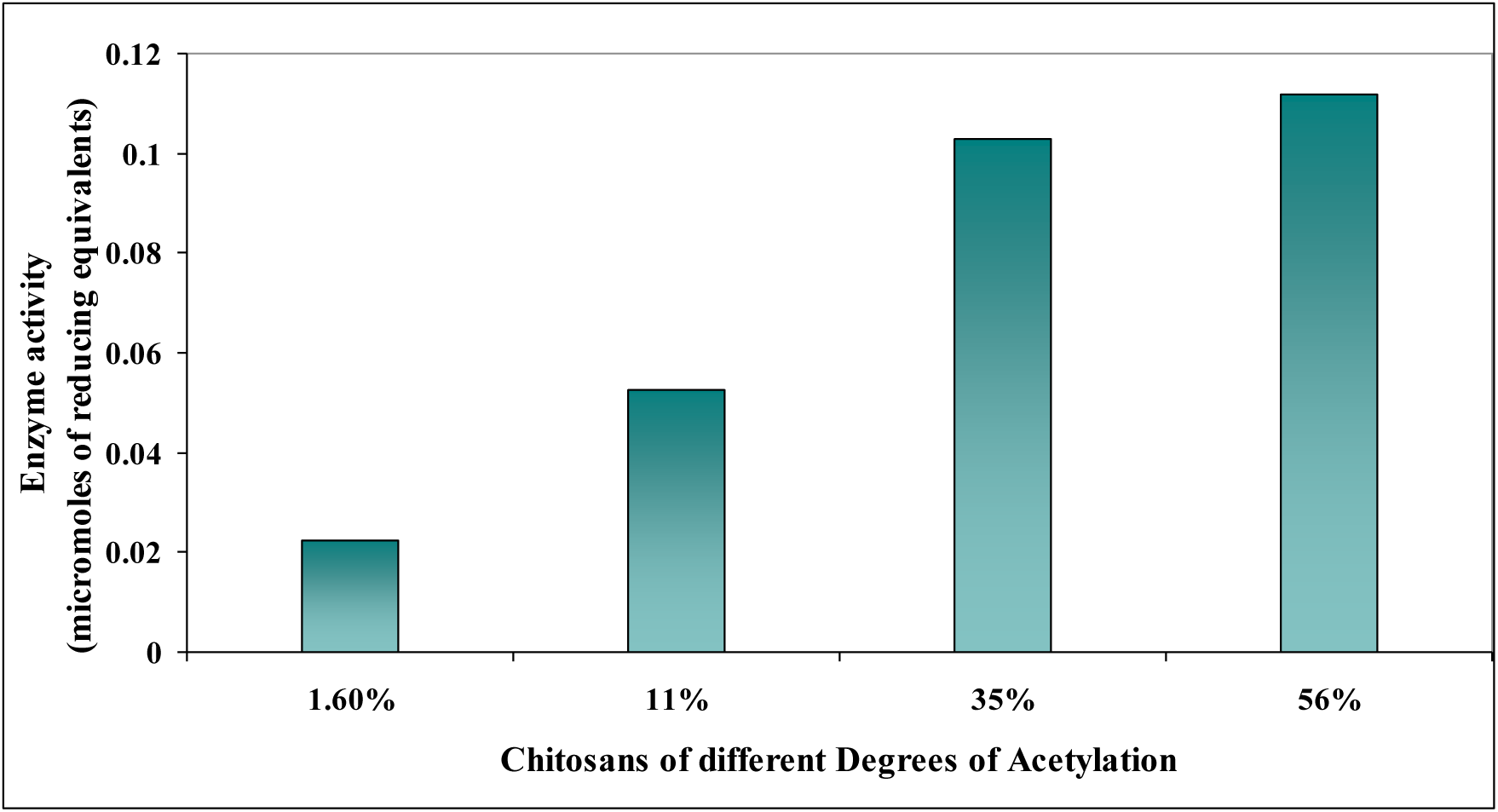
Effect of chitosans of different degrees of acetylation on *P. aeruginosa* P17 CCME activity

##### Km and Vmax of the enzyme

The inverse values of substrate concentration effect on enzyme activity were used to arrive at the Lineweaver-Burk plot (double reciprocal plot of enzyme kinetics) and using the linear equation of the plot, Km and Vmax values of the P17 chitinase were calculated and they are 2.38 uM and 8.33 umole/ min respectively.

##### Hydrolysis of chitosans

Chitinase of P17 showed differential hydrolysis pattern of low and high DA chitosans. The enzyme could not digest 11% DA chitosan after 24 h whereas DA 56% chitosan was digested to some extent and released oligomers *viz* di-, tri- and tetra-mers. However, after 48 h partial digestion of 11% DA chitosan was observed with the release of mono- and di-mers and comparatively higher digestion of 56% DA chitosan was seen with the release of a few di- to hexa-mers (Fig 13).

**Fig 13:**
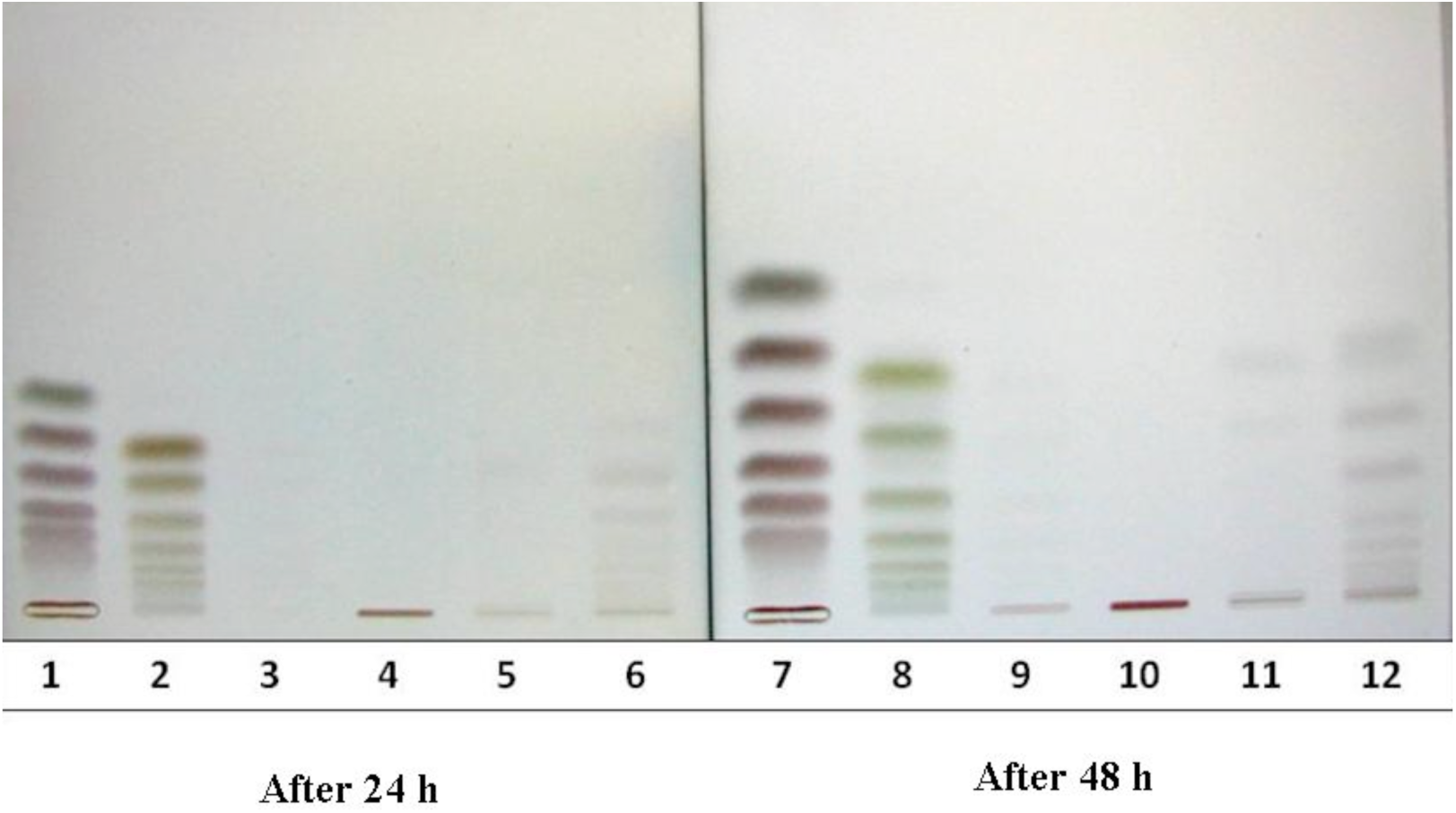
Thin layer chromatogram showing the digestion products of low and high DA chitosans using *P. aeruginosa* P17 CCME {Lane 1, 2 & 7, 8 are chitin/chitosan standards (di-mer to hexa-mers); Lane 3, 4 & 9, 10 are DA 11 and 56% controls; Lane 5, 6 & 11, 12 are enzyme digested chitosans}

In another experiment completely acetylated (A2, A3, A4, A5 and A6) oligomers of chitin and de-acetylated (D2, D3, D4, D5 and D6) oligomers of chitosan were used as substrates to understand the enzyme activity. The chitinases of P17 could not digest both A2 and D2. A3 was digested releasing di- and mono-mers whereas, A4, A5 and A6 were hydrolyzed into tri-, di- and mono-mers. P17 CCME could not hydrolyze D3, D4 and D5 (Fig 14).

**Fig 14:**
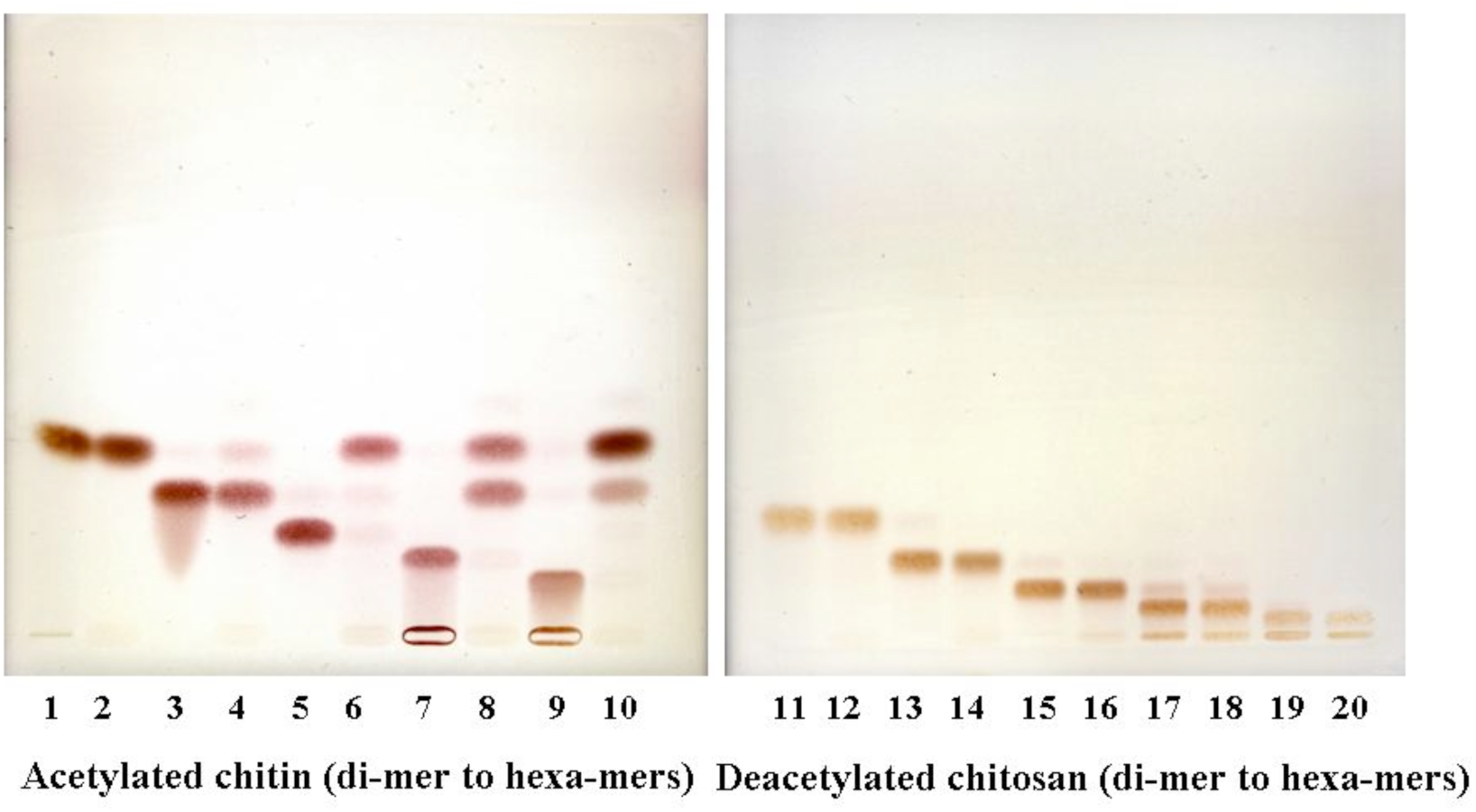
Thin layer chromatogram showing the digestion products of di- to hexa-mers of chitin (A) and chitosan (D) after 24 h {Lane with odd numbers are di-mer to hexa-mer (A and D) controls; Lane with even numbers are enzyme treated samples}

##### Substrate digestion and oligomer release

Time-course digestion monitoring by high performance size exclusion chromatography showed that 90% of the chitosan substrate was digested to oligomers in the very first hour. Then the released oligomers were further digested increasing the concentration of tetra- and penta-mers which have a molar mass of less than 1×10^3^ gmol^−1^ (Fig 15).

**Fig 15:**
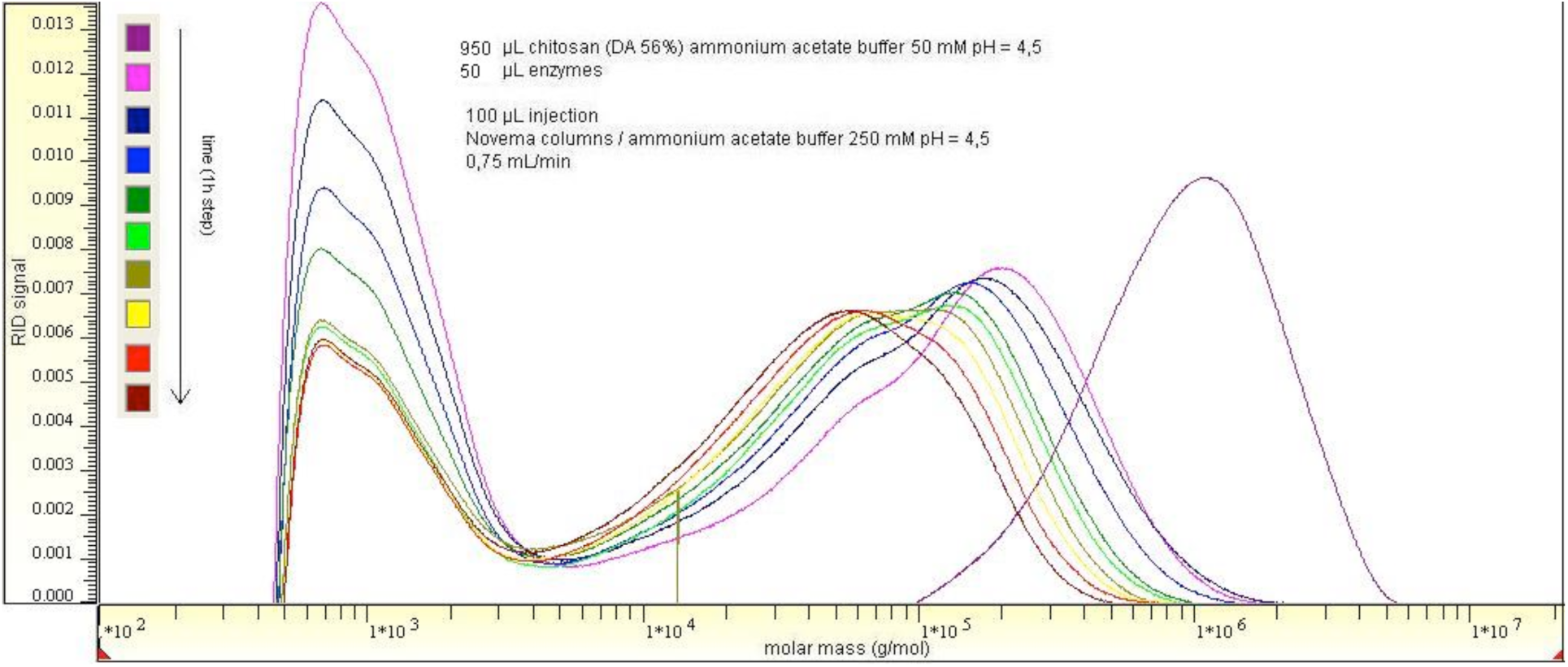
Size exclusion chromatogram showing reduction in molar mass of chitosan DA 56% with *P. aeruginosa* P17 chitinase enzyme

##### Detection of oligomer size by MALDI-o-TOF

Different sized oligomers ranging from 2-mer to 11-mer were detected and the sequence of the same was deduced. Result showed that the minimum sized oligomer produced was an acetylated dimer (A2 with m/z of 447.15) with low abundancy. Different oligomers observed were, trimer (D1A2; m/z 608.21), tetramer (D2A2 with m/z 769.3 & D1A3 with m/z 811.29), pentamer (D2A3 with m/z 1133.4), hexamer (D2A4 with m/z 1175.4, D3A3 with m/z 1225.5), heptamer (D4A3 with m/z 1294.5, D3A4 with m/z 1336.5, D2A5 with m/z 1378.5), octamer (D4A4 with m/z 1497.6, D3A5 with m/z 1539.6) and followed by 9 sized products (D5A4 with m/z 1658.6, D4A5 with m/z 1700.7), 10 sized (D6A4 with m/z 1819.7, D5A5 with m/z 1861.8) and finally 11 sized (D6A5 with m/z 2022.8, D5A6 with m/z 2226.9) (Fig 4.35). Of all the detected oligomers D2A2, D2A3, D3A3, D3A4 and D6A4 abundancy was high (Fig 16).

**Fig 16:**
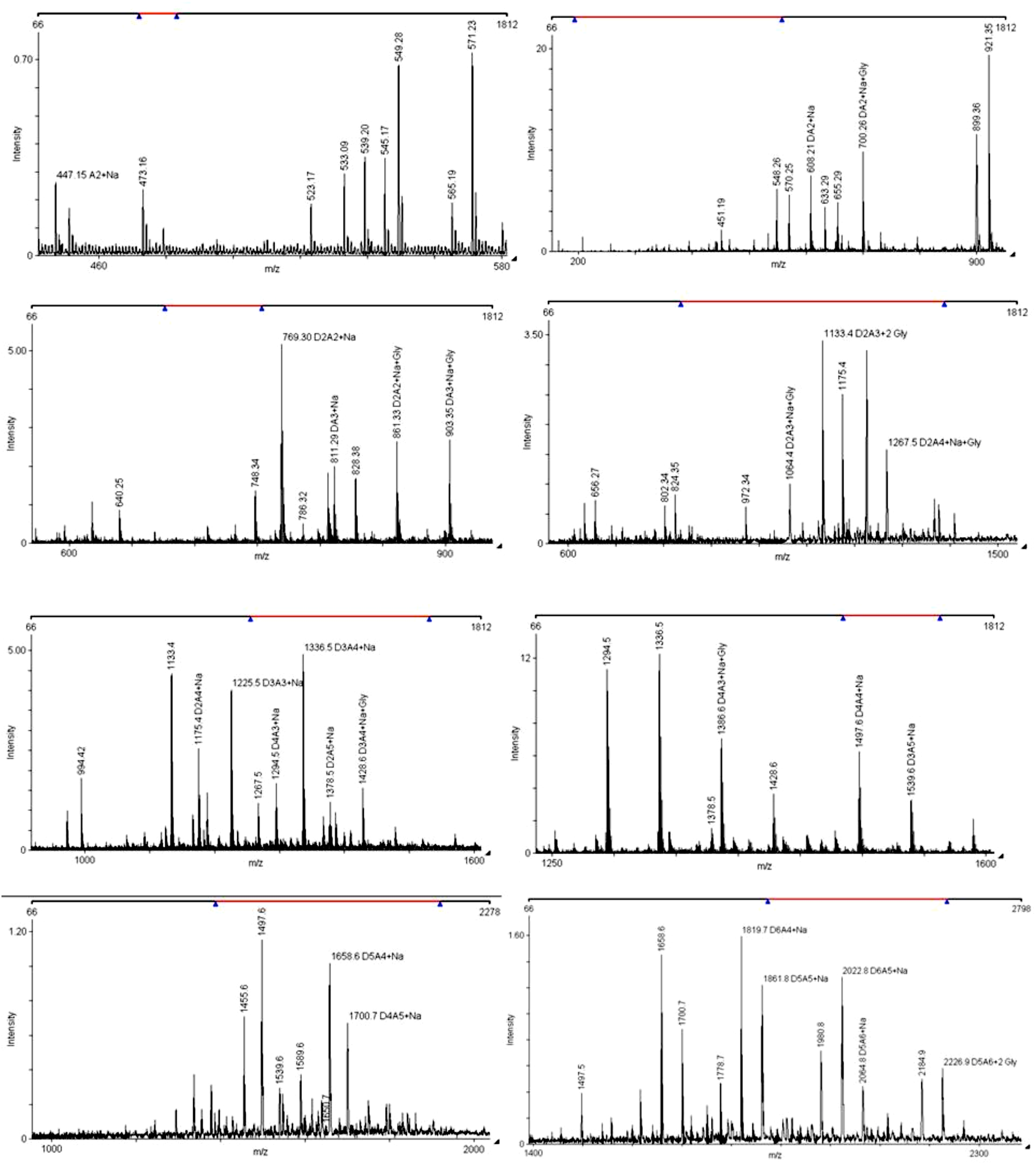
Mass signals (MALDI-o-TOF) showing different oligomers produced after digestion of chitosan DA 56% by *P. aeruginosa* P17 chitinase

##### Chitinase identification by 2D-PAGE

About ten (10) different peptides were identified such as, and the peptide sequences showed homology with the known chitinase of *P. aeruginosa* from the database. The spot on activity gel after calcoflour white staining was used for chemical characterization by mass spectrometry. Different peptides were identified as shown in Table 5; and Fig 17.

**Fig 17:**
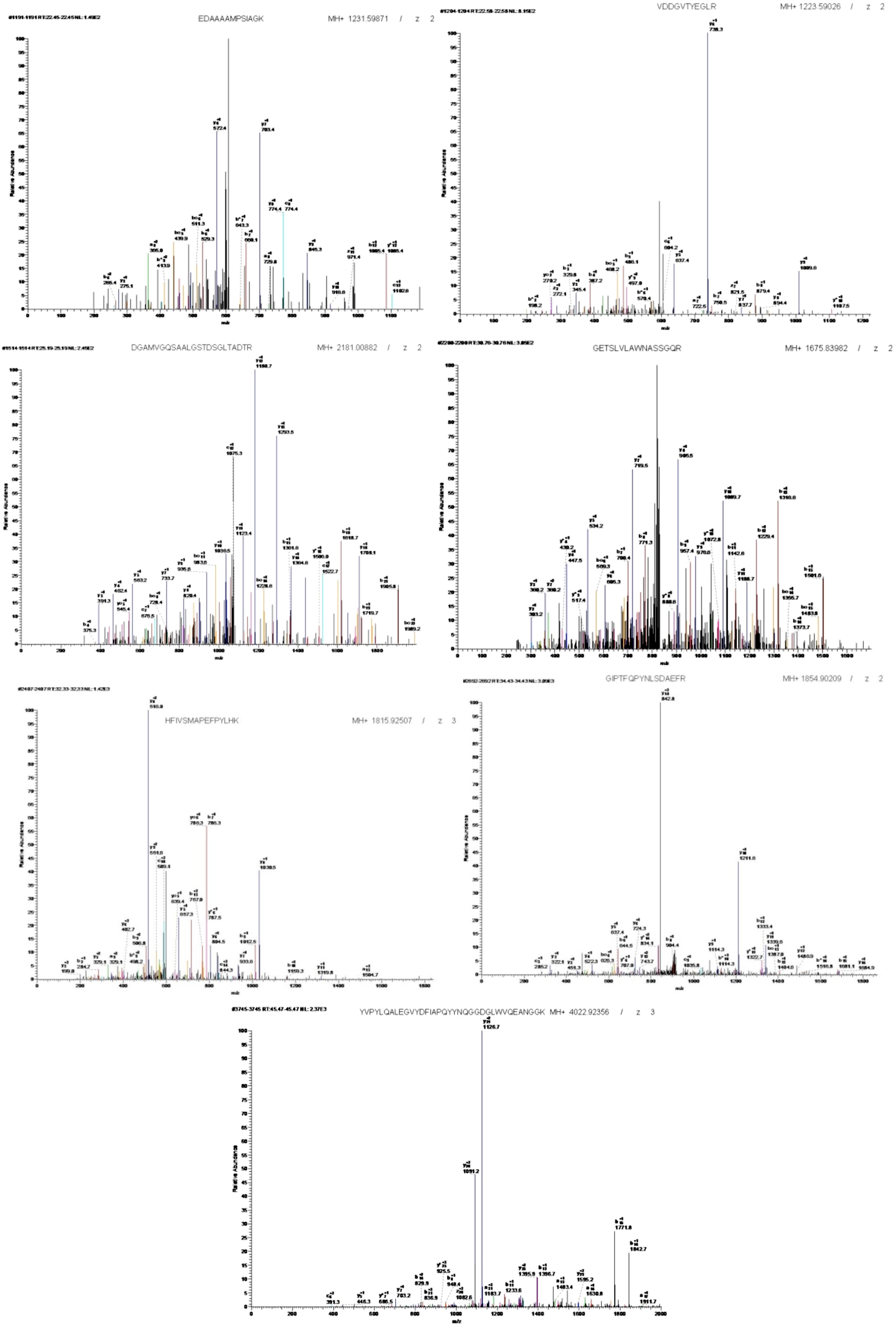
Mass spectrum analysis showing different peptide sequences of *P. aeruginosa* P17 chitinase

**Table 5:**
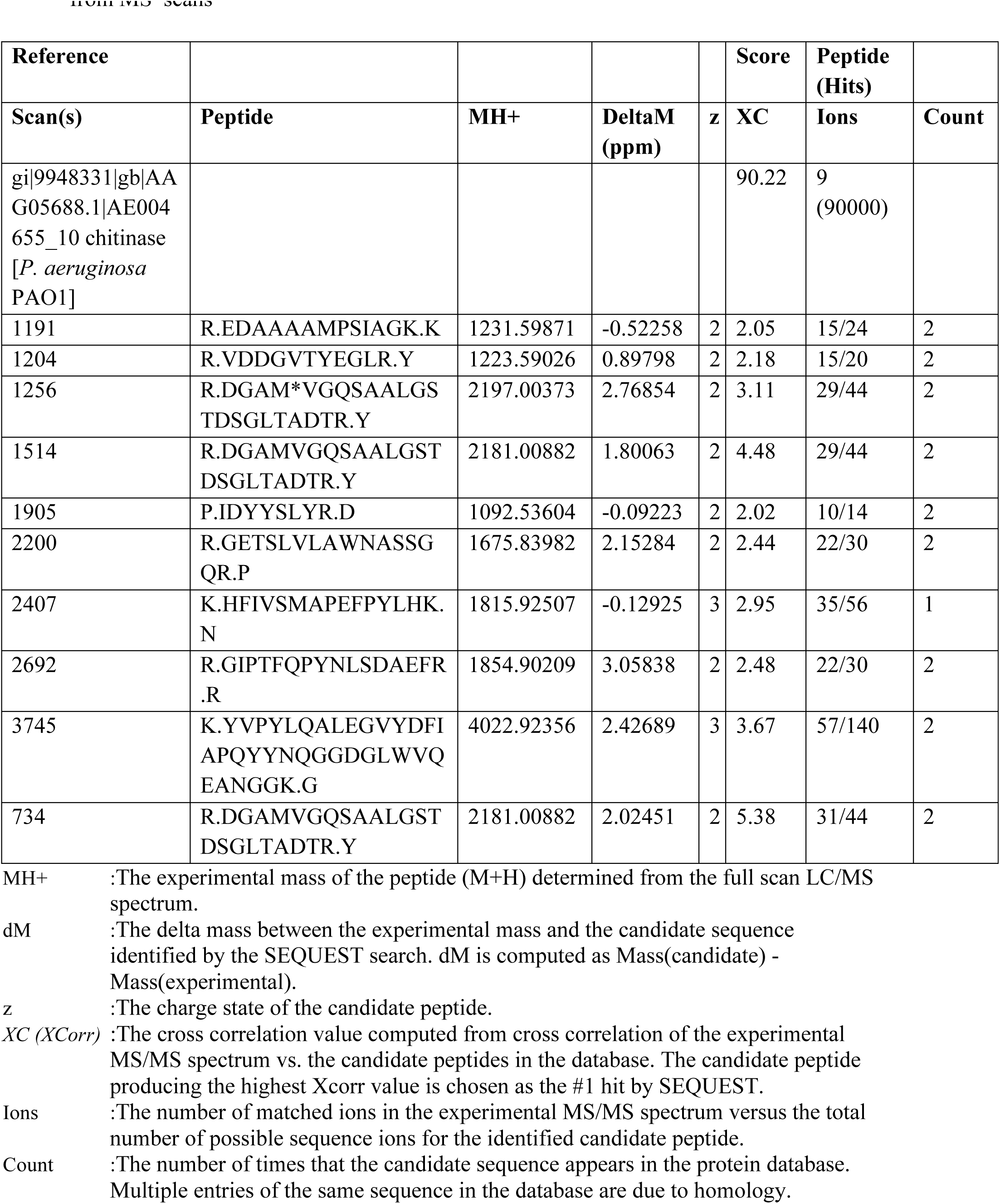
Characteristics of peptide (*P. aeruginosa* P17 chitinase) sequences obtained from MS scans

## 4. Discussion

Four (04) strains of Pseudomonas spp. namely P1, P17, P22 and P28 were selected for the present experiments on sorghum due to their ability to promote growth of sorghum and pigeon pea under in vitro and glass house conditions and possessing abiotic stress tolerance and also have noted plant growth promoting traits which has been previously demonstrated and evidence with publications (data not shown). All the strains were identified up to species level using ribotyping and biochemical characterization (Table 1).

### 4.1 Effect of bacterial-chitosans consortium on sorghum growth

With the aim of developing *Pseudomonas*-chitosan combination for plant growth promotion, bacterial growth response was studied by incubating them in the presence of low and high degree of acetylation (DA) chitosans. It was observed that in presence of 1.6% DA chitosan (with higher positive charge) the growth of all four selected strains of Pseudomonas significantly reduced as compared to 56% DA chitosan. This could be due to the fact that the bacteria are completely cross linked by the high positive charge of DA 1.6% resulting in reduced growth. On the other hand, with high DA chitosan, due to less number of positive charges on chitosan, higher growth was observed (Fig 1). The antibacterial effect of chito-oligosaccharides has been shown to be greatly dependent on their degree of polymerization (DP) or molecular weight and requires glucosamine polymers with DP 6 or greater (Kendra and Hadwiger 1984). Since P1 has no demonstrated chitinolytic activity this was used as a negative control check for the current experiments.

In the current study, chitosan treated plants showed higher growth compared to that of control plants (Table 4). The plant growth promoting effect of chitosan may be due to its root strengthening and stem thickening properties. Plant growth promoting effect of radiation degraded chitosans was demonstrated by Luan et al. (2002) where radiation degraded products of DA 80% chitosan enhanced root length, shoot length and biomass of *Limonium latifolium, Eustoma grandiflorum* and *Chrysanthemum morifolium* flower plants.

In present study also all the treatments showed higher growth in terms of root length, shoot length and dry mass (Table 4). DA 56% chitosan treated plants recorded more dry mass. However, the growth of plants treated with bacteria was higher than chitosan alone treated plants. This might be due to PGPR activity of the microbe leading to higher availability of nutrients in the rhizosphere, production of phytohormones and expression of various PGPR traits. Among the individual bacterial treatments, P17 treated plants recorded highest root length, shoot length and dry mass. Among the combined treatments, chitosan DA 56% in combination with bacteria showed higher growth than chitosan DA 1.6% + bacterial treated plants. This could be due to lower growth of bacteria in presence of DA 1.4% and thus lesser expression of PGPR traits. Further, among DA 56% + bacterial combinations, chitosan DA56% in presence of P17 showed highest growth probably due to the PGPR activity of this strain. Root length of these plants was more and this could be due to IAA secretion by P17 coupled with enhanced rooting of plants in presence of chitosan (Table 4 and Fig 2-3). Chitinolytic PGPR have been reported to enhance the growth of banana crop under field conditions (Kavinoa et al. 2010).

Many studies have shown effect of chitosans and chitinolytic bacteria suppressing plant pathogens. Plant growth promoting and disease suppressing ability of chitosans was showed in rice plant by Suchada et al. (2008), whereas spraying of chitosan solution remarkably enhanced some pigments in tomato (El-Tantawy 2009). These reports suggest that chitosan which contains about 8.7% nitrogen might promote both vegetative and reproductive growth of some plants.

Plants when exposed to chitosans usually activate their defence system followed by expression of various defence enzymes and pathogenesis related (PR) proteins. So, in the current study it was planned to estimate these defence enzymes which could play an important role in induced systemic resistance. Chitinases induced by PGPR play an important role in PGPR mediated insect management by hydrolyzing the chitin, as chitin constitutes a structural component of the gut linings of insects (Harish et al. 2009).

In the present study, all the defence enzymes increased significantly in chitosan treated and chitosan-bacterial treated plants. Higher PAL activity was shown by plants treated with Pseudomonas P22+chitosan DA 56% combination. The presence of chitosan and chitosan degraded products in the root region of the plants might have made plants to accumulate more PAL enzyme (Fig 4). Similar results were also shown in experiments conducted on soybean (Wajahatullah Khan et al. 2003). Since PAL is a key enzyme in the phenylpropanoid pathway, its activity lead to synthesis of phenols and compounds that were associated with expression of resistance (Nicholson and Hammerschmidt 1992). Chitosan causes biological effects as plant growth promotion, the direct growth inhibition of several microorganisms, mainly fungi and elicits induced resistance in plants against their pathogens (Bautista-Baños et al. 2006). In all chitosan treated plants, enzymes like peroxidase and poly phenol oxidase increased (Fig 5-6) which might be due to the elicitor activity of chitosans and oligomers obtained by chitosan digestion in the rhizosphere region of plants. Phenolics content of plants was also enhanced significantly in chitosan treated plants (Fig 7).

Recent investigations on mechanisms of biological control by PGPRs revealed that some PGPR strains protect plants from various pathogens by inducing resistance in plants. In the present study also the combination of Pseudomonas and chitosan enhanced various defence enzymes in plants than their individual counterparts suggesting that this combination could be more effectively employed for plant growth and to enhance resistance in plants. Even the two different pathogenesis related proteins like chitinase and glucanase levels increased significantly upon treatment of chitosan and bacteria in sorghum plants indicating their protective role (Fig 8-9). Of all the treatments Pseudomonas P17 + chitosan DA 56% plants accumulated high levels of most of the defence enzymes. Hence this combination could be recommended for promotion of growth of sorghum and ISR in plants. Chitinase and peroxidase, have been reported as the most important components of the induced systemic resistance (Nandakumar et al. 2000).

### 4.2 Characterization of CCME in strain P17

In the current study, chitin-chitosan modifying enzymes (CCME) of *Pseudomonas aeruginosa* P17 was characterized because of its plant growth promoting ability towards selected crops and hydrolytic activity on wide range of chitosans. CCME of P17 had one active polypeptide with a Pi in the range of 3.0-4.0 and the activity staining also confirmed this by showing only one thick band on gel containing chitosan DA 56% (Fig 10-11). The molecular weights of most microbial chitinases are in the range of 40–65 kDa. The presence of one isoform of chitinase suggested that the enzyme functionality was governed by only one protein.

Digestion of low and high DA chitosans was studied to know the various hydrolytic products produced by the enzyme. After 48 h interval, di-, tri-, tetra-, penta- and hexa-mers were released as a result of hydrolysis of chitosan. Further the enzyme could not digest acetylated (A) and deacetylated (D) dimers where as tri-, tetra-, penta- and hexa-mers were digested indicating that probably the enzyme has endochitinase activity (Fig 13-14) and it required at least a tri-mer to show the activity. Similar results were also reported by Moon-Sun Jang et al. (2005).

Substrate specificity assay showed that the enzyme had more specificity towards highly acetylated chitosans (Fig 12) indicating that probably it has more chitinase activity and less chitosanase activity. Higher reducing ends were estimated when enzyme was incubated with chitosan DA 56% substrate and the number of reducing ends released was less with decrease in DA of the substrate confirming that the enzyme has more specificity for highly acetylated chitosans.

Biological control with fluorescent pseudomonads offers an effective method of managing plant pathogens (Ramamoorthy et al., 2001). In the present study, the appearance of a clearing zone indicated that P. aeruginosa P17 isolate could degrade chitin substrate. Bacterial chitinases are known to show a broad range of iso-electric points with 4.5-8.5 (Watanabe et al, 1990), where the isolate in current study showed a pI in the range of 3.0-4.0. Only, one iso-form of the enzyme could be detected followed by dot blot of iso-electric focusing of gel. This could be an advantage that the enzyme does not need any other proteins to be functional in nature, compared to other chitinases as reported by Wang et al. (2008) who reported that two different enzymes of Pseudomonas sp. TKU015 are required for chitin/ chitosan digestion. Multiple isozymes of endochitinase from *Pseudomonas fluorescens* were demonstrated by Nielsen and Sorensen (1999). Wang et al. reported five extracellular chitinases of Bacillus cereus 6E1 using an in-gel chitinase assay. Substrate specificity study showed that the enzyme was more specific to high DA substrates than low DA substrates. This explains that the enzyme has high chitinase and low chitosanase activities. This was further evidenced by the observation of thin layer chromatogram where hydrolysis of 11 and 56% DA chitosans was identified. These results are in agreement with the findings of Moon-Sun Jang et al. (2005). The enzyme of P17 isolate could not digest dimers of either A or D form which further denotes that it requires a minimum of trimer in order to show activity. Therefore, it gives preliminary information that the enzyme has ‘endo’-chitinase kind of activity.

Degradation of chitosan-56% DA substrate with Pseudomonas-P17 sp. chitinase enzyme followed by separation of the hydrolysis products using size exclusion chromatography revealed that more than 90% of the substrate was digested during the initial first hour. Processive degradation is thought to improve catalytic efficiency because single polymer chains are prevented from re-associating with the insoluble material in between catalytic cycles (20, 21), thus reducing the number of times the enzyme has to carry out the energetically unfavorable process of gaining access to a single chain. These results suggest that the apparent link between processivity and enzyme efficiency observed for chitinases is a more general phenomenon, which may be valid for many enzyme-substrate systems. The knowledge on the role of processivity of the enzyme gives ideas for the development of enzymatic tools for biomass conversion.

Because of the crystalline and inaccessible nature of chitin and to some extent, chitosan enzymes could have developed special tactics to ensure efficient degradation and this could be the reason in the current study that the enzyme has shown certain degree of processivity. It is also known from studies that chitin-degrading organisms produce accessory proteins that disrupt the crystalline substrate, thus increase the efficiency of chitinases (Karkehabadi et al. 2008). The detection of oligomers by MALDI-o-TOF showed the least sized oligomer was an acetylated dimer and the digestion pattern revealed that the enzyme always needed the presence of atleast one ‘A’ group in the polymer to show the hydrolysis (Fig 7). The low molecular weight oligomers released could also play an important role in inhibition of pathogenic organisms in soil as they have antimicrobial action. Yalpani et al. (1992) found that chitosan oligomers with varying degrees of polymerization reduced the growth of E. coli. Various sized chitosan oligomers produced by hydrolysis of chitin play a key role in antibiosis (Sang-Hoon and Samuel, 2004). Identification of the 2D spot followed by tryptic digestion revealed the presence of 10 different peptides that could be correlated to the protein of *Pseudomonas aeruginosa* chitinase PA01 strain of GenBank (accession no. 9948331).

## 5. Conclusions

In conclusion, unlike most of the other reports regarding chitinase producing strains of *Pseudomonas* sp., this research aimed at the partial characterization, processivity of the enzyme and identification of chitinase from P. aeruginosa-P17 strain and development of Pseudomonas (P17) and Chitosan 56% DA consortium for enhanced plant growth and induced systemic resistance in plants. Further studies regarding the detailed mechanism of enzyme action, and other kinetics related parameters need to be studied for developing improved biocontrol agents and biological-chemical tailor made products for agriculture use.

## Acknowledgements

Duetscher Akademischer Austausch Dienst (DAAD)-German Academic Exchange Service sandwich fellowship to Praveen Kumar, G. is gratefully acknowledged.

